# The live attenuated *DGAT1*-knockout whole-cell *Toxoplasma* vaccine confers protective immunity against acute and chronic toxoplasmosis

**DOI:** 10.64898/2026.08.16.745102

**Authors:** Shahbaz M. Khan, Yevel Flores-Garcia, Jiro Sakai, Mustafa Akkoyunlu, Julia D. Romano, Karen Ehrenman, Viviana Pszenny, Michael E. Grigg, Krishna S. Manuguri, Yue Zhao, Duanpei Wang, Fidel Zavala, Isabelle Coppens

## Abstract

The intravacuolar parasite *Toxoplasma gondii* scavenges fatty acids from host mammalian cells and stores excess in lipid droplets. To investigate the physiological relevance of neutral lipid storage in *Toxoplasma*, we generated a mutant lacking *DGAT1*, an ER-localized enzyme that synthesizes triacylglycerols, from the virulent type I RH strain of *T. gondii*. Compared to WT, RH ΔDGAT1 parasites grow poorly in mammalian cells, form few LD, suffer from lipotoxicity, and do not cause disease or lethality in immunocompetent or immunodeficient mice. Importantly, mice immunized with RH ΔDGAT1 parasites mount strong, long-term immune responses involving both cellular and humoral components, with higher levels of *T*. *gondii*-specific IgG antibodies, effector memory T cells, and both pro-inflammatory and anti-inflammatory cytokines, indicating a mixed Th1/Th2 response with Th1 predominance. This immunity provides complete, long-lasting protection (up to 6 months) against rechallenge from homologous type I (acute infection) and heterologous cyst-forming type II (chronic infection) *T*. *gondii* strains. Additional analyses reveal that IFN-γ, CD8^+^ T cells, as well as B cells are crucial for defending against type I *T*. *gondii* in immunized mice. Overall, our live-attenuated RH ΔDGAT1 strain is a promising vaccine candidate and a model for studying immune responses that control *T*. *gondii* infections.

## Introduction

*Toxoplasma gondii* is an obligate intracellular apicomplexan parasite that forms infectious tissue cysts in the brain and muscles of various animal hosts. It causes toxoplasmosis, a zoonotic infection mainly transmitted by contact with contaminated Felidae feces (oocysts containing sporozoites). Warm-blooded animals, including humans, typically become infected through multiple transmission routes: ingesting oocysts from the environment (via contaminated water or soil) or tissue cysts containing bradyzoites (via contaminated muscle meat), or through congenital transmission from infected mothers to their offspring (1). Oocysts are shed in large amounts in infected cat feces, remain extremely stable in the environment, resist most inactivation methods, and are highly infectious (2). Due to the easiness and high prevalence of infection, with one in three people globally being chronically infected with *T*. *gondii* (1), CDC has classified toxoplasmosis among the five neglected parasitic infections (3). Research in the mid to late 2000s identified toxoplasmosis as the second leading cause of deaths from foodborne illnesses and the fourth leading cause of hospitalizations due to foodborne illnesses in the United States (4). In humans, this infection causes severe brain and eye disease and permanent damage to the fetus during pregnancy or in situations of immunosuppression (1). In small ruminants, *T*. *gondii* infection primarily leads to abortion and stillbirth, causing heavy production losses (5).

Treatment in humans typically combines the antifolate drugs pyrimethamine and sulfadiazine, which were first shown to be effective in mice more than 70 years ago (6). However, hypersensitivity and toxicity limit the efficacy of this treatment (7). This primary therapy for toxoplasmosis targets tachyzoites (proliferative forms derived from sporozoites or bradyzoites) associated with acute illness but is ineffective against dormant bradyzoites found in tissue cysts, allowing recrudescence of active infections that may lead to severe complications (7, 8). Moreover, despite considerable efforts, no safe and effective drug is available for treating toxoplasmosis in farm ruminants (9). An effective vaccine should provide rapid, long-lasting immunity in vaccinated individuals and minimize disease in livestock with a reduction in shedding of oocysts in felines, thereby preventing the spread of the disease. However, despite extensive efforts and knowledge about the immune response to *T*. *gondii*, no vaccine exists for human toxoplasmosis (10). The only approved vaccine in the world is Toxovax^®^, a live-attenuated *T. gondii* S48 strain intended for farm ruminants but does not provide complete efficacy. This live-attenuated vaccine has an unknown genetic background, raising safety concerns (11). As such, there is a pressing need to develop safe and effective vaccines with identified correlates of protection to address this public health concern. Recently, various vaccine candidates reviewed in (10, 12)—including inactivated, recombinant protein subunit, DNA, mRNA, exosome, carbohydrate, nanoparticle, and live-attenuated vaccines—have been tested across multiple animal models, including sheep and goats. So far, these candidate vaccine strategies have not succeeded in eliminating tissue cysts or preventing congenital transmission. Among these platforms, live-attenuated vaccines are considered the most effective at inducing immune protection because they expose the immune system to a wide range of immunogens (10, 13).

Lipids are energy-dense molecules and serve as essential building blocks for membrane biosynthesis in animals, plants, and fungi. Cytosolic lipid droplets (LD) are active organelles that store neutral lipids, support metabolic needs, and protect against lipotoxicity and membrane damage in eukaryotic cells. In mammals, the membrane-bound *O*-acyltransferase (MBOAT) family includes the acyl-CoA:cholesterol acyltransferases (ACAT1/2) and the acyl-CoA:diacylglycerol acyltransferase 1 (DGAT1) enzymes, which acylate cholesterol and diacylglycerol to form the major neutral lipids, cholesteryl ester (CE) and triacylglycerol (TAG), respectively (14). *Toxoplasma* expresses TgDGAT1, an important MBOAT that detoxifies energy-rich fatty acids and stores them as TAG in intracellular LD (15). We have previously shown that T863, a DGAT1 inhibitor originally developed to treat obesity in humans, can impair parasite growth without harming host cells (16). However, the direct impact of DGAT1 deletion in *Toxoplasma* has not yet been thoroughly investigated.

In this study, we demonstrate that deleting the *DGAT1* gene from the highly virulent *Toxoplasma* type I RH strain results in LD depletion, limiting the parasite’s ability to survive intracellularly. We also show that the RH ΔDGAT1 strain is avirulent in both immunocompetent and immunodeficient mice, even at high lethal doses. This weakened infection triggers a robust humoral and cellular *Toxoplasma*-protective immune response in mice, as demonstrated by immunodepletion studies during rechallenge with homologous type I and heterologous cyst-forming type II strains, respectively. Our results show that the non-cyst-forming RH ΔDGAT1 strain is a promising vaccine candidate against toxoplasmosis and a helpful tool to define protective immune responses against *Toxoplasma* infections.

## Results

### The growth of RH ΔDGAT1 strain is significantly attenuated and shows reduced LD biogenesis

Previous attempts to disrupt *DGAT1* from the *Toxoplasma* genome (TGGT1_232730 in www.ToxoDB/org) by homologous recombination-based knockout strategies were unsuccessful, suggesting that the TgDGAT1 gene is essential for parasite survival (15). However, using the highly accurate CRISPR/Cas9 genome-editing technology that possesses increased homologous recombination efficiency (Supplementary Figure 1a) and a modified cloning strategy, we successfully targeted the deletion of *TgDGAT1* by replacing its gene by an HXGPRT selection cassette in the *Ku80*-deleted type I RH strain (17). After multiple rounds of selection, single knockout clones were obtained, but all exhibited significant growth defects, compared to parental (control) parasites. The deletion of *DGAT1* and insertion of the HXGPRT selectable marker in mutant clones were confirmed at the genomic level by multiple diagnostic PCR using deletion– and integration-specific primers (Supplementary Figure 1b and c).

To compare the replication ability of RH ΔTgDGAT1 (hereafter called KO) parasites with that of the parental (hereafter called WT) parasites in fibroblasts, we first measured their replication rates at 24 hours post-infection (p.i.) by counting parasites per parasitophorous vacuole (PV), as illustrated by immunostaining using antibodies against the parasite plasma membrane marker SAG1. Most PVs in the WT group contained 4 to 8 parasites, whereas most KO PVs had only 1 to 2 parasites that had lost the characteristic crescent shape of *T*. *gondii* (Figure 1a-c). By 48 hours, the difference in replication rates between WT and KO parasites became more significant. We also found that at least 11% of KO strain PVs had “odd” parasite counts whereas the WT strain did not produce any such aberrant PV (Figure 1d and e), indicating abnormal, asynchronous replication within the PV. Plaque assays were conducted to evaluate the KO parasites’ invasion ability (plaque counts) and their growth (plaque areas) over several cycles of combined invasion, replication and egress. At day 10 p.i., the KO strain produced notably fewer and smaller plaques in fibroblasts compared to the WT strain, suggesting defects in invasion and growth, thus a considerable loss of fitness due to an impaired lytic cycle (Figure 1f). Several attempts to complement the KO strain by inserting a C-terminal V5-tagged DGAT1 coding sequence driven by an endogenous *T*. *gondii* promoter either into the UPRT gene locus (18) or into a neutral locus suitable for transgene integration (19), using CRISPR-Cas9 gene editing, did not yield viable parasites. This suggests that the KO strain lacks sufficient robustness for electroporation-based gene editing.

**Fig. 1.**
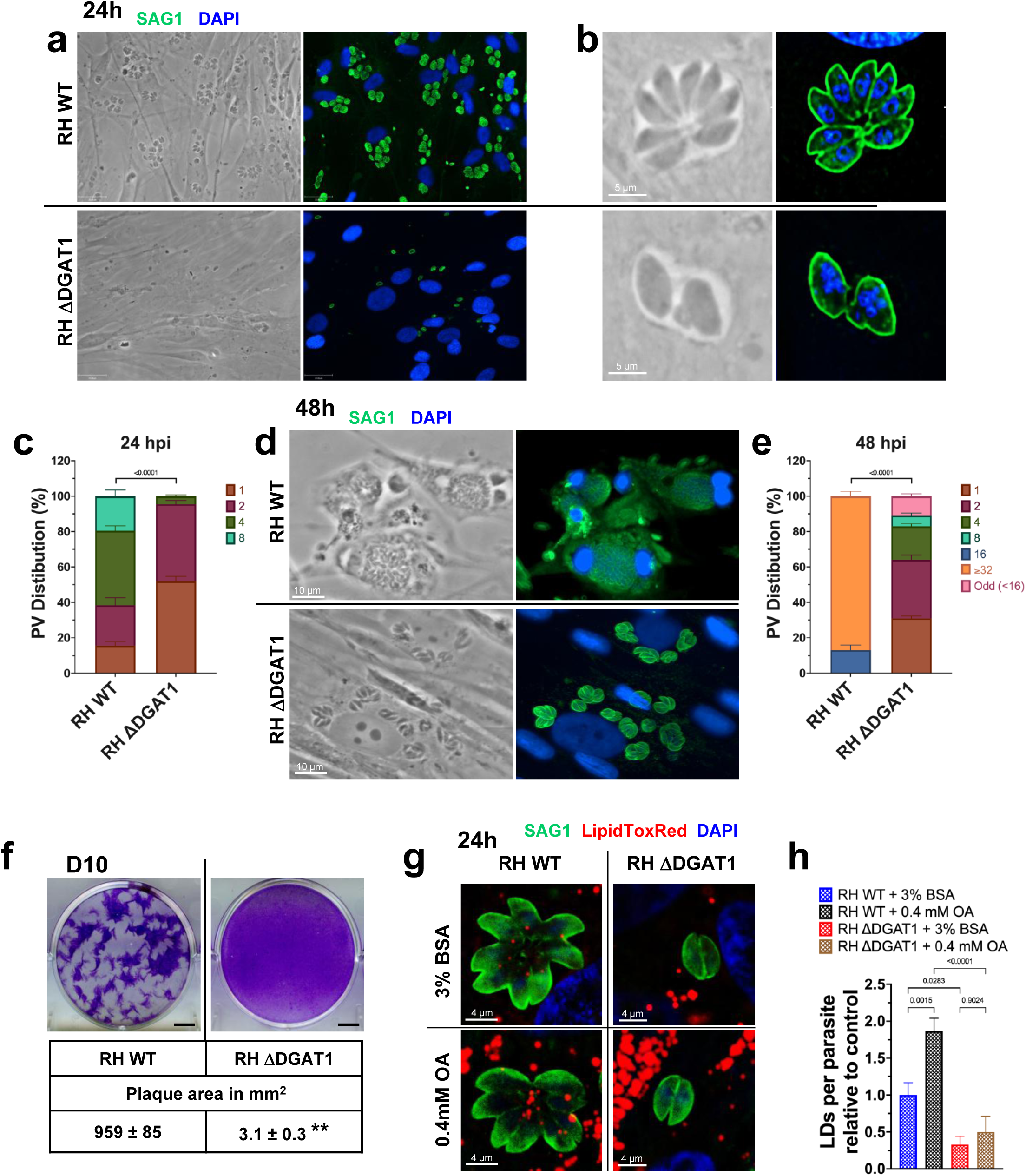
Phenotypic analyses of the ΔDGAT1 *Toxoplasma* mutant reveal severe growth and replication defects. (a-c) Immunofluorescence microscopy of RH WT– or RH ΔDGAT1-infected fibroblasts at 24 h post-infection shows (a) smaller PVs (reduced replication) with (b) misshapen parasites in ΔDGAT1-infected fibroblasts. Infected human foreskin fibroblasts (HFFs) were fixed with 4% formaldehyde and stained with anti-SAG1 antibody (plasma membrane marker) and DAPI. (c) Mutant parasites exhibited delayed replication, resulting in reduced replication efficiency, as indicated by the distribution of parasites within PVs at 24 h post-infection. (d) At 48 h post-infection, fluorescence microscopy of infected fibroblasts shows smaller ΔDGAT1 PVs in contrast to very large WT PVs close to egress, as confirmed by (e) replication quantification. Both experiments were performed twice, each with two replicates, and at least 100 PVs were counted per sample. The mean ± SD data were analyzed using two-way ANOVA. The “Odd” group includes PVs in which the number of parasites was either 3, 5, 6, 7, or 12. The representative images shown at the 24 h and 48 h time points are from two independent experiments. (f) Plaque assays comparing parental and ΔDGAT1 strains revealed that ΔDGAT1 mutants have defects in invasion and intracellular growth. Fibroblast monolayers were infected with 150 RH WT or RH ΔDGAT1 parasites, and their invasion and growth were assessed after 10 days by counting and measuring plaque sizes. Representative images and quantification of the lysed area (means ± SD) from 3 independent experiments are shown; **, *p* < 0.0025 (unpaired two-tailed t-test). Scale bars, 5.25 mm (g-h) TgDGAT1 has an important role in LD biogenesis. (g) RH WT– or RH ΔDGAT1-infected HFFs under control conditions (0.3% BSA) or excess fatty acids (0.4 mM OA complexed to 0.3% BSA) were fixed at 24 h after infection with 4% formaldehyde and stained with anti-SAG1 antibody, LipidToxRed (LD marker), and DAPI. (h) Imaging analysis was performed to visualize the LipidToxRed signal for LD detection within the parasite, revealing increased signal in WT parasites under excess OA, in contrast to mutant parasites. Representative images and data (mean ± SD) from 2 independent experiments are shown. A one-way ANOVA followed by Tukey’s multiple comparisons test was used to evaluate statistical significance.

We next investigated the role of TgDGAT1 in LD biogenesis in *T*. *gondii* by LD counting in WT and KO parasites at 24 hours p.i. (Figure 1g and h). Deleting *DGAT1* significantly reduced LD biogenesis in *T*. *gondii*. Exogenous addition of oleate markedly increased LD biogenesis in the WT strain but did not significantly stimulate LD production in the mutant strain, indicating that TgDGAT1 is an important contributor to LD formation in *T*. *gondii*.

To scrutinize the ultrastructure of KO parasites, we performed electron microscopy (EM) analyses at day 4 and 10 p.i. *Toxoplasma* divides by endodyogeny, a form of asexual reproduction in which two identical daughter cells are assembled within the mother cell, with organelles either recycled from the mother cell or formed de novo, resulting in the synchronous emergence of the progeny (20). A striking phenotype of the KO at day 4 was the loss of temporal regulation in daughter cell formation and mother cell consumption, resulting in misshapen parasites (Figure 2Aa), consistent with our immunofluorescence assays (IFA) in Figure 1b. Another observation was the accumulation of abnormal electron-dense, lipophilic material within the PV, whose origin is unknown, but this suggests lipid disorders associated with excess membrane accumulation (Figure 2Aab). Two hallmarks of *Toxoplasma* infection are the close apposition of host mitochondria and ER to the PVM (21) and the formation of an intravacuolar network (IVN) (22) that traps nutrient-rich organelles (23), and these features were observed in the KO. However, the PV lumen of the mutant contained many uncharacterized, round structures, more abundant at day 10 p.i. (Figure 2Aa and Bab). At that time, KO parasites exhibited more membrane defects, including protrusions and bulges on the parasite surface (Figure 2Bc) or deep multilayered invaginations (Figure 2Bd). Many IVN tubules were observed appended to the PVM, suggesting maintained scavenging activities (Figure 2Be).

### The RH ΔTgDGAT1 mutant is avirulent in immunocompetent as well as immunodeficient mice

The WT RH strain exhibits high virulence in all laboratory mouse strains, with an estimated 100% lethal dose of a single viable parasite (24). To evaluate the in vivo virulence of DGAT1-deficient *T*. *gondii*, we tested the knockout (KO) strain in a BALB/c mouse model of lethal toxoplasmosis (Figure 3a). Mice infected with the WT strain experienced acute infection, rapidly lost weight, and died within 8 days, whereas mice infected with the KO strain displayed no symptoms and survived beyond 30 days (Figure 3b and c). We also challenged interferon-gamma knockout (IFN-γ KO) mice because IFN-γ is critically important in host resistance to *T. gondii* infection as mice lacking IFN-γ quickly die from toxoplasmosis, even when infected with avirulent strains (25). As expected, infecting IFN-γ KO mice with WT strain led to rapid weight loss and quick death within 10 days. In contrast, even at a high lethal dose, the KO strain remained avirulent in IFN-γ KO for over 30 days, demonstrating a significant reduction in virulence (Figure 3d and e). During *Toxoplasma* infection, NK and T cells, especially CD8^+^ T cells are the main producers of IFN– γ (26–28). To interrogate the role of these cells in host resistance to WT and KO strains, we challenged triple-immunodeficient NCG mice that lack functional/mature T, B, and NK cells and have reduced macrophage and dendritic cell functions. Resembling the susceptibility profile of IFN-γ KO mice, NCG mice rapidly lost weight and succumbed to challenge with WT strain by day 8 (Figure 3d and g). As observed in IFN-γ KO mice, not only the KO strain infected NCG mice survived the challenge but also progressively gained weight.

**Fig. 2.**
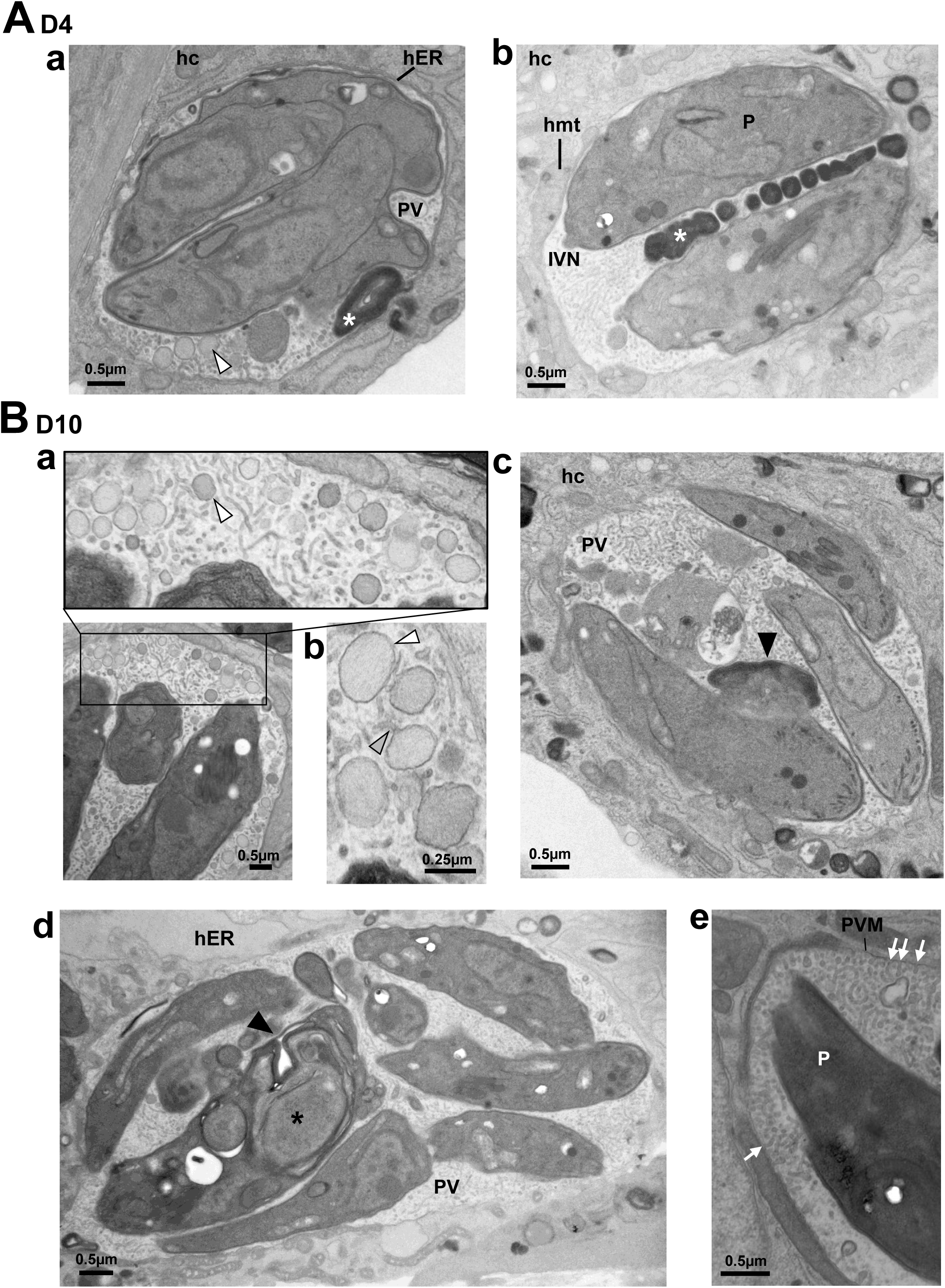
Ultrastructure of DGAT1-deficient *Toxoplasma* strain reveals cytopathies with membrane abnormalities Human fibroblasts were infected with RH ΔDGAT1 parasites for 4 (A) or 10 days (B), then fixed and processed for EM. In A: representative EM images of ΔDGAT1 showing a PV with abnormally dividing parasites (panel a) or a PV with two daughters (panel b). The PVM is associated with the host ER and mitochondria, and the PV lumen is filled with the IVN, along with abnormal electron-dense lipophilic structures (white asterisks) and grayish round structures (white arrowhead) of unknown origin. (c-g) In B: representative EM images of ΔDGAT1 illustrating the intra-PV accumulation of the round structures (white arrowheads) connected to the IVN tubules (gray arrowhead) (panels a and b), mutant parasites with membrane damage (black arrowheads in panels c and d), and a PV containing many IVN tubules appended to the PVM (white arrows in panel e). hc, host cell; hER, host endoplasmic reticulum; hmt, host mitochondria; IVN, intravacuolar network; PV, parasitophorous vacuole; P, parasite; PVM, PV membrane.

**Fig. 3.**
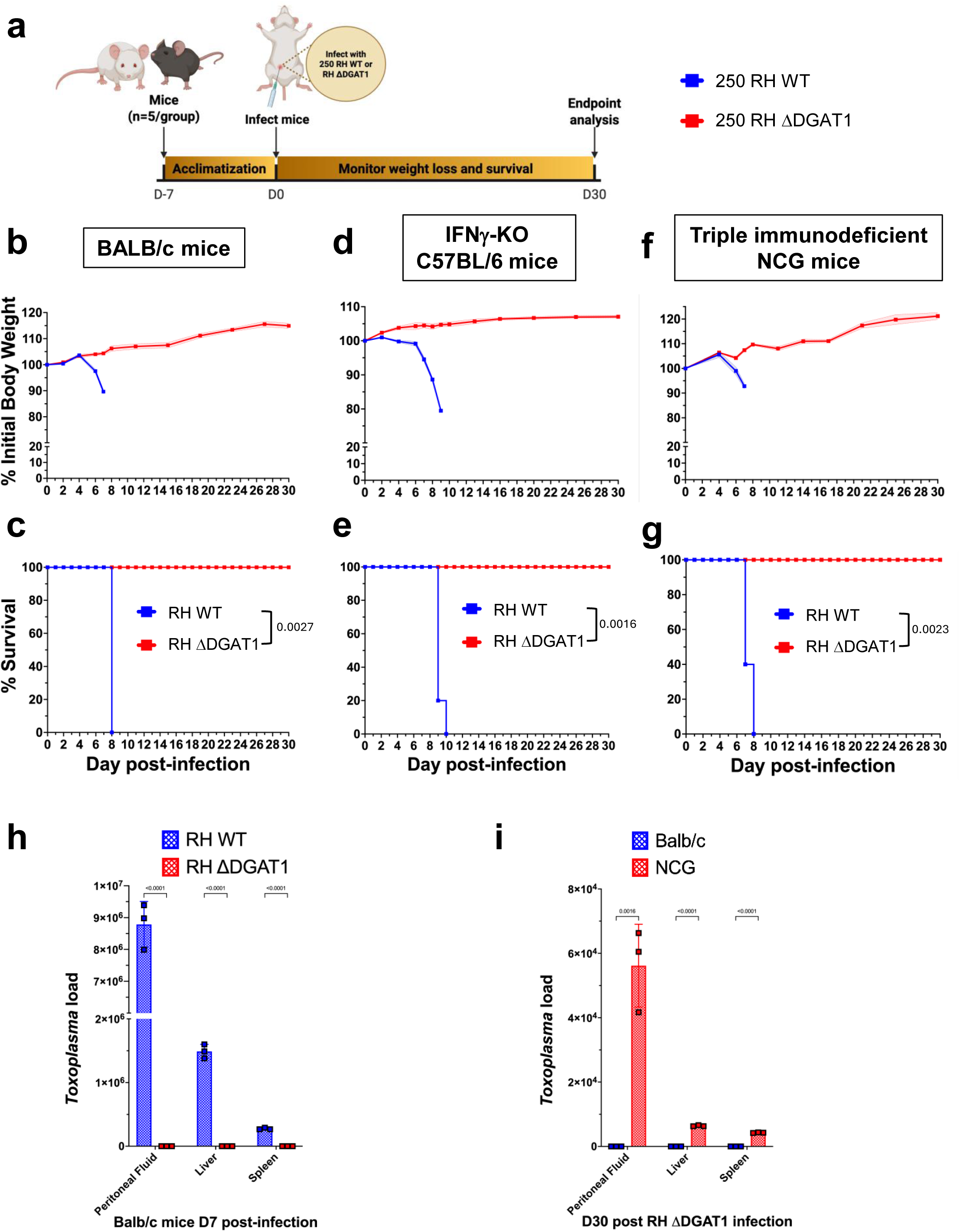
Ablation of TgDGAT1 abolishes the virulence of *T*. *gondii* in immunocompetent and immunodeficient mice. (a) Freshly egressed tachyzoites (250) from either the RH WT or RH ΔDGAT1 *T*. *gondii* strain were injected intraperitoneally into BALB/c, IFN-γ KO, or triple immunodeficient NCG mice, which lack T cells, B cells, and NK cells (n=5 mice per group for each strain). The body weight and survival of the infected mice were monitored over 30 days. The mutant strain did not cause body weight loss or death in (b, c) BALB/c mice, (d, e) IFN-γ KO mice, or (f, g) NCG mice. Mean ± SEM body weight data are shown. Statistical significance of the probability of survival was assessed using the log-rank (Mantel–Cox) test. (h) Parasite burdens were undetectable in immunocompetent mice infected with the ΔDGAT1 mutant at day 7 post-infection. BALB/c mice were injected with 250 WT or ΔDGAT1 tachyzoites, and parasite loads in the peritoneal fluid, liver, and spleen of mice were detected 7 days post-infection by qPCR (mean ± SD data from 3 biological replicates and analyzed by multiple unpaired two-tailed t-tests). (i) By day 30 after infection, parasite levels were undetectable in immunocompetent mice, while only barely detectable in mice with triple immunodeficiency. BALB/c and NCG mice were infected with 250 RH ΔDGAT1 tachyzoites, and parasite burden in the peritoneal fluid, liver, and spleen of mice was detected 30 days post-infection by qPCR (mean ± SD data from 3 biological replicates and analyzed by unpaired two-tailed t-test).

Additionally, qPCR analysis of parasite burden in the peritoneal fluid, liver, and spleen of BALB/c mice at 7 days p.i. revealed undetectable parasite levels in KO-infected mice, compared to WT-infected mice that exhibited very high parasite loads (*P*<0.0001) in these tissues (Figure 3h). When testing KO-infected immunocompetent BALB/c and immunodeficient NCG mice at day 30 of infection, parasite DNA was not detected in BALB/c mice, although a low level of parasite load was observed in healthy-looking NCG mice at that time (Figure 3i).

### RH ΔTgDGAT1 inoculation in mice induces a strong humoral immune response

The complete avirulence of the RHΔDGAT1 strain in immunodeficient mice prompted us to investigate its potential to induce a robust *T*. *gondii*-specific immune response. However, achieving this required a sufficient number of parasites for vaccination purposes. Therefore, we aimed to improve the growth of KO parasites in cell culture by adjusting culture conditions. Our lab typically cultivates *Toxoplasma* parasites in minimum essential medium (MEM) supplemented with 10% FBS and a physiologically relevant 1 g/L glucose concentration. However, the mutant ΔDGAT1 strain showed poor growth in this medium, as evidenced by less uracil incorporated into the parasites and reduced size of lysis plaques (Supplementary Figure 1a and b). To address this, we used MEM supplemented with 10% lipid-depleted FBS (LPD-FBS) and increased the glucose concentration to 4.5 g/L to improve the mutant strain’s growth. We anticipated that this adjustment would shield the mutated parasites from free fatty acid (FFA) toxicity, allowing them to utilize glucose as an alternative energy source instead of lipids. Indeed, the mutant parasites showed significantly better growth in the modified medium.

We inoculated inbred BALB/c mice by intraperitoneal injection and noted that an inoculum of either 10^4^ or 10^5^ KO parasite doses did not cause a long-term reduction in the body weight of mice (data not shown). To examine the antibody responses induced by RH ΔDGAT1 inoculation, we analyzed the production of *Toxoplasma*-specific total IgG and IgG1/IgG2a subtypes in serum samples from naïve and inoculated mice at various time points up to day 180 post-infection using ELISA (Figure 4a). We observed high OD values for IgG at all time points in inoculated mouse groups compared to the uninoculated group (*P* < 0.0001), at a serum dilution of 1/2,700 (Figure 4b and d). Furthermore, the data indicated a dose-dependent response. Mice inoculated with 10^5^ RH ΔDGAT1 tachyzoites exhibited higher IgG production compared to those inoculated with 10^4^ RH ΔDGAT1 tachyzoites. Notably, at any given serum dilution, IgG1 OD values were lower than the IgG2a ones, pointing that the inoculation may elicit a Th1-biased immune response in mice (Supplementary Figure 3). Interestingly, the parasite dose influenced the Th1/Th2 nature of the immune response, with a low 10^4^ tachyzoite dose tending to promote predominantly a Th1 response, while a higher 10^5^ tachyzoite dose favored a mixed Th1/Th2 response at day 30 post-inoculation (Figure 4e). By day 180, however, both groups stabilized into a Th1-dominated, mixed Th1/Th2 response through Th1/Th2 switching.

**Fig. 4.**
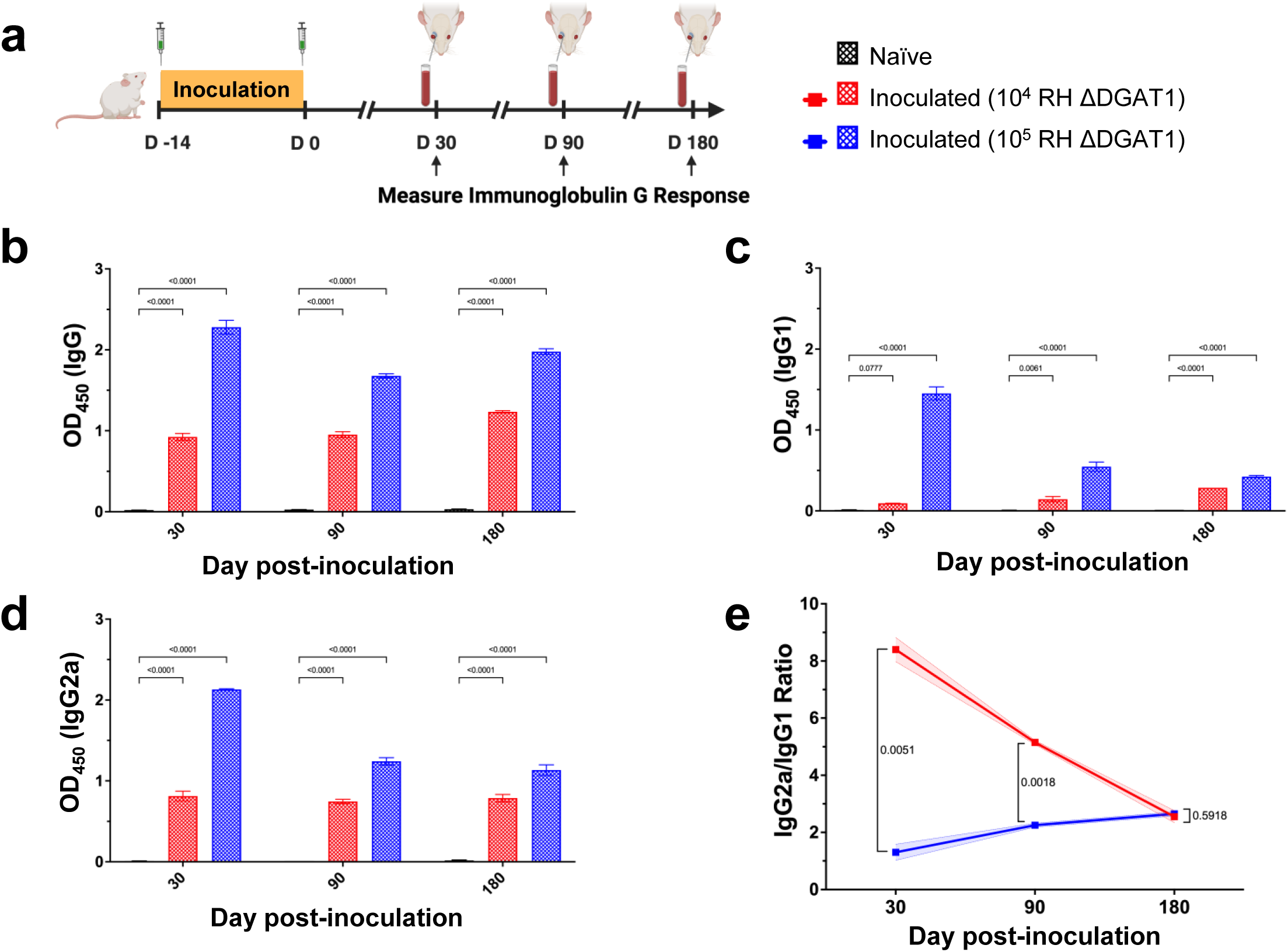
Vaccination using the ΔDGAT1 mutant elicits robust antibody-mediated immune responses against *T. gondii* infection in mice. (a) Timeline for detection of *T. gondii*-specific IgG response in BALB/c mice by ELISA. Mice (n = 5 per group) were inoculated with two doses of 10^4^ or 10^5^ RH ΔDGAT1 parasites, and IgG, IgG1, and IgG2a levels, and the IgG2a/IgG1 ratio, were measured in the serum of naïve and vaccinated mice at 30, 90, and 180 days post-inoculation. Relative levels of *Toxoplasma*-specific IgG were determined by indirect ELISA using immuno-plates coated with 10 μg/mL of soluble *T*. *gondii* antigens. (b-e) Data for a serum dilution of 1/2700 are shown. Mice immunized with RH ΔDGAT1 showed significantly higher levels of (b) IgG and its subclasses (c) IgG1 and (d) IgG2a than the control group (mean ± SD data analyzed by two-way ANOVA with Dunnett’s multiple-comparison test). (e) The IgG2a/IgG1 ratio at 30 days post-vaccination indicated that the vaccine dose influences the Th1/Th2 balance: a lower dose tends to promote a predominantly cellular (Th1) response, whereas a higher dose favors a mixed Th1/Th2 response. This pattern was not observed at 180 days post-vaccination. Statistical analyses were performed using multiple unpaired t-tests with Holm-Sidak’s multiple-comparison test. Mean ± SD data with *p*-values are shown.

### Mouse inoculation with RH ΔDGAT1 parasites triggers a robust cellular immune response

To assess the cell-mediated immune response following RH ΔDGAT1 inoculation, we measured the proportions of effector memory CD4^+^ and CD8^+^ T lymphocytes in the peripheral blood of naïve and inoculated mice using the activation marker CD44 and the main homing receptor CD62L (29–31). Specifically, we analysed the percentages of long-lived CD4^+^CD44^+^CD62L^-^ and CD8^+^CD44^+^CD62L^-^ memory T cells among peripheral CD3^+^ T lymphocytes by flow cytometry six months after RH ΔDGAT1 vaccination (Figure 5a). We noted that blood collected from inoculated mice contained a significantly higher, dose-dependent proportion of CD4^+^ and CD8^+^ T cells with the cell-surface phenotype CD44^+^CD62L^-^, characteristic of memory cells, compared with that in uninoculated mice. (Figure 5b and c). Our data suggest that KO inoculation elicits CD4^+^ and CD8^+^ T lymphocytes with a memory phenotype with likely potential to recall immunity against *T*. *gondii*.

**Fig. 5.**
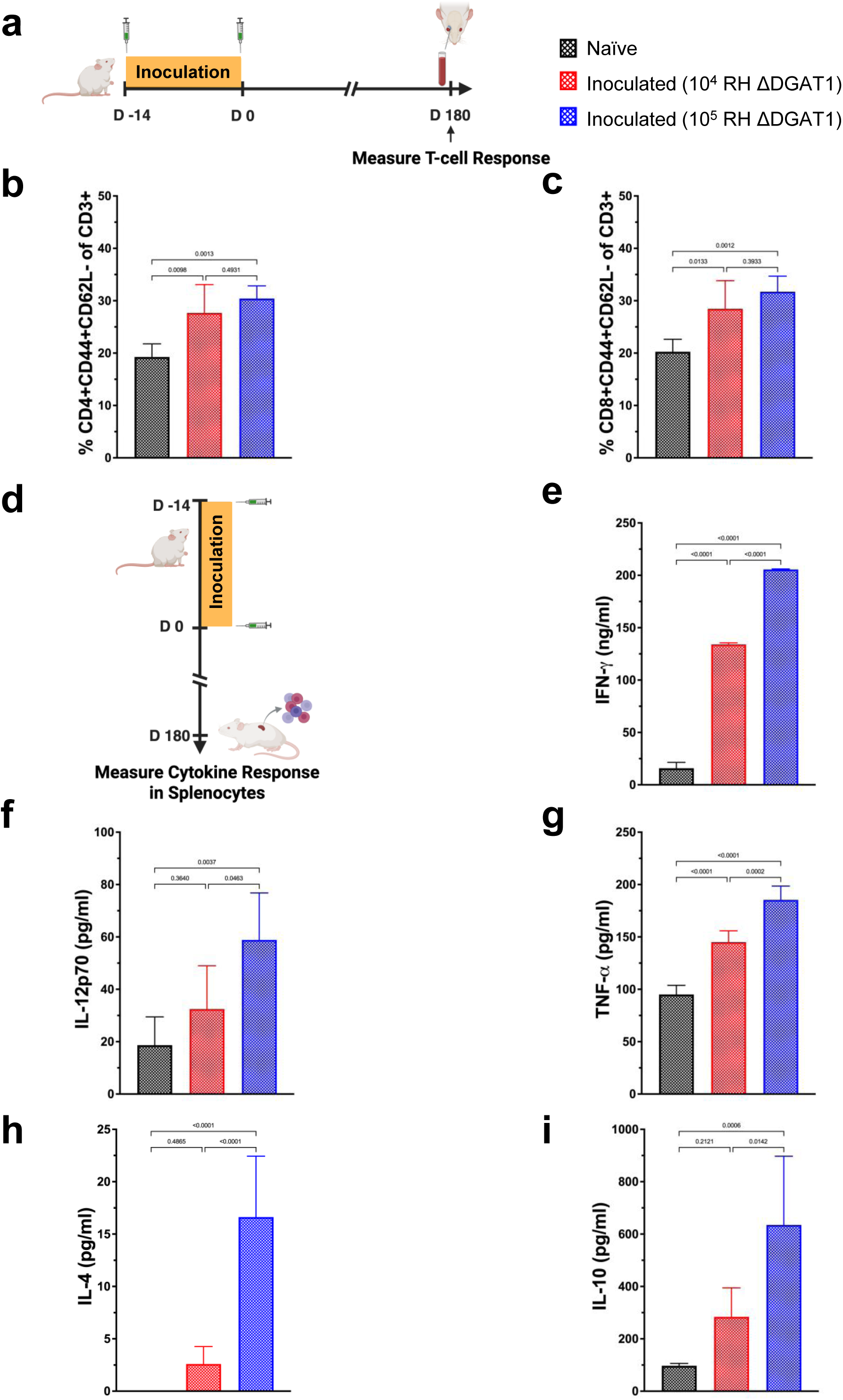
Inoculation of the ΔDGAT1 mutant induces strong cellular immune responses against *T. gondii* infection in mice. (a) Timeline for detection of the memory T-cell response in BALB/c mice by flow cytometry. The proportions of CD44^+^CD62L^-^ effector memory CD4^+^ and CD8^+^ T cells within peripheral CD3+ T lymphocytes in the blood of vaccinated mice were measured at 180 days post-inoculation. Lymphocytes from non-vaccinated mice served as negative controls. Vaccinated mice exhibited a notable increase in long-lived (b) CD4^+^ and (c) CD8^+^ memory cells compared to the control group. Data are expressed as a percentage of total CD3^+^ T cells and presented as means ± SD (n = 5 mice per group). Statistical significance was assessed using one-way ANOVA with Tukey’s multiple-comparison test. (d) Timeline for measuring the cytokine response by splenocytes from BALB/c mice by ELISA. (e-i) Cytokine (IFN-γ, IL-12p70, TNF-α, IL-4, and IL-10) production by splenocytes harvested from immunized and naïve mice was assessed at 180 days post-immunization with RH ΔDGAT1 (n = 5 mice per group). Splenocytes were stimulated in vitro with soluble *T. gondii* antigens (10 μg/mL). Cell-free supernatants were collected and tested for IL-4 activity after 24 hours, for TNF-α and IL-10 activity after 72 hours, and for IL-12p70 and IFN-γ activity after 96 hours. Mice immunized with RH ΔDGAT1 showed significantly stronger pro-inflammatory (e, IFN-γ; f, IL-12p70; g, TNF-α) and anti-inflammatory (h, IL-4; i, IL-10) cytokine production compared to the control group. Bars represent the means ± SD, with indicated *p*-values determined by ordinary one-way ANOVA with Tukey’s multiple-comparison test.

We measured pro-inflammatory cytokines (IFN-γ, IL-12p70, TNF-α) and anti-inflammatory cytokines (IL-4, IL-10) in supernatants from splenocyte cultures of naïve and inoculated mice after 180 days (Figure 5d). These cultures were stimulated with soluble *Toxoplasma* antigens and tested using ELISA. Cytokine levels were measured at different times: IL-4 at 24 hours, TNF-α and IL-10 at 72 hours, and IL-12p70 and IFN-γ at 96 hours post-stimulation. Our findings show that inoculation with RH ΔDGAT1 parasites led to a significant increase in both pro-inflammatory and anti-inflammatory cytokines, which remained elevated even after 180 days, compared with uninoculated controls, indicating a long-term memory response (Figure 5e-i). Cytokine production also exhibited a dose-dependent response, consistent with *Toxoplasma*-specific IgG levels in serum samples and memory T cells in the spleens. In general, mice inoculated with 10^5^ RH ΔDGAT1 tachyzoites elicited a stronger cytokine response than those inoculated with 10^4^ RH ΔDGAT1 tachyzoites. However, this trend was more pronounced for anti-inflammatory cytokines than for pro-inflammatory cytokines: the group inoculated with 10^5^ tachyzoites exhibited significantly higher levels of Th2-type cytokines, particularly IL-4, compared to uninoculated mice, more so than the group inoculated with 10^4^ tachyzoites (Figure 5h and i). This indicates that mice receiving higher KO doses exhibit more robust activation of anti-inflammatory mechanisms and that their immune response gradually shifts toward a balanced Th1/Th2 profile over time. Among all cytokines tested, IFN-γ exhibited the highest levels, significantly surpassing the others, highlighting its crucial role as the primary activator of cell-mediated immunity for *T*. *gondii* clearance.

### Inoculation with RH ΔTgDGAT1 parasite confers long-lasting protection against both type I and type II *Toxoplasma* infections in mice

To evaluate the potential of the RH ΔDGAT1 strain as a whole-tachyzoite genetically attenuated vaccine against acute toxoplasmosis, we infected outbred CD-1 mice with 10^3^ to 10^5^ KO parasites by intraperitoneal injection. We challenged naïve and inoculated mice at 30, 90, or 180 days with a lethal intraperitoneal dose of 500 luciferase-expressing tachyzoites from the homologous RH Type I strain (RH-Luc) (Figure 6a and f; Supplementary Figure 4a). Inoculation with RH ΔDGAT1 parasites protected mice from an acute, lethal challenge of RH parasites, regardless of the inoculate dose or the timing of challenge. Inoculated mice exhibited no signs of infection and gained weight over the 30-day monitoring period. In contrast, naïve mice lost weight rapidly and died within 11-12 days of the acute challenge (Figure 6b, c, g and h; Supplementary Figure 4b and c). Bioluminescence imaging revealed severe infection in naïve mice by days 5 or 6 after the challenge, worsening by day 9, while all inoculated mice were infection-free on day 21 post-challenge (Figure 6d, e, i and j; Supplementary Figure 4d and e). We also evaluated immunity in mice inoculated with mutant parasites that survived the WT challenge after 30 days (Supplementary Figure 4f). These vaccinated mice were rechallenged with WT RH parasites after 90 days and were protected against acute lethal toxoplasmosis, with no body weight loss or death during the monitoring period (Supplementary Figure 4g and h). In vivo imaging also established that mice were free of WT RH parasites by day 21 and never reached the level of infection displayed in naïve mice by parasite-derived luminescence (Supplementary Figure 4i and j).

**Fig. 6.**
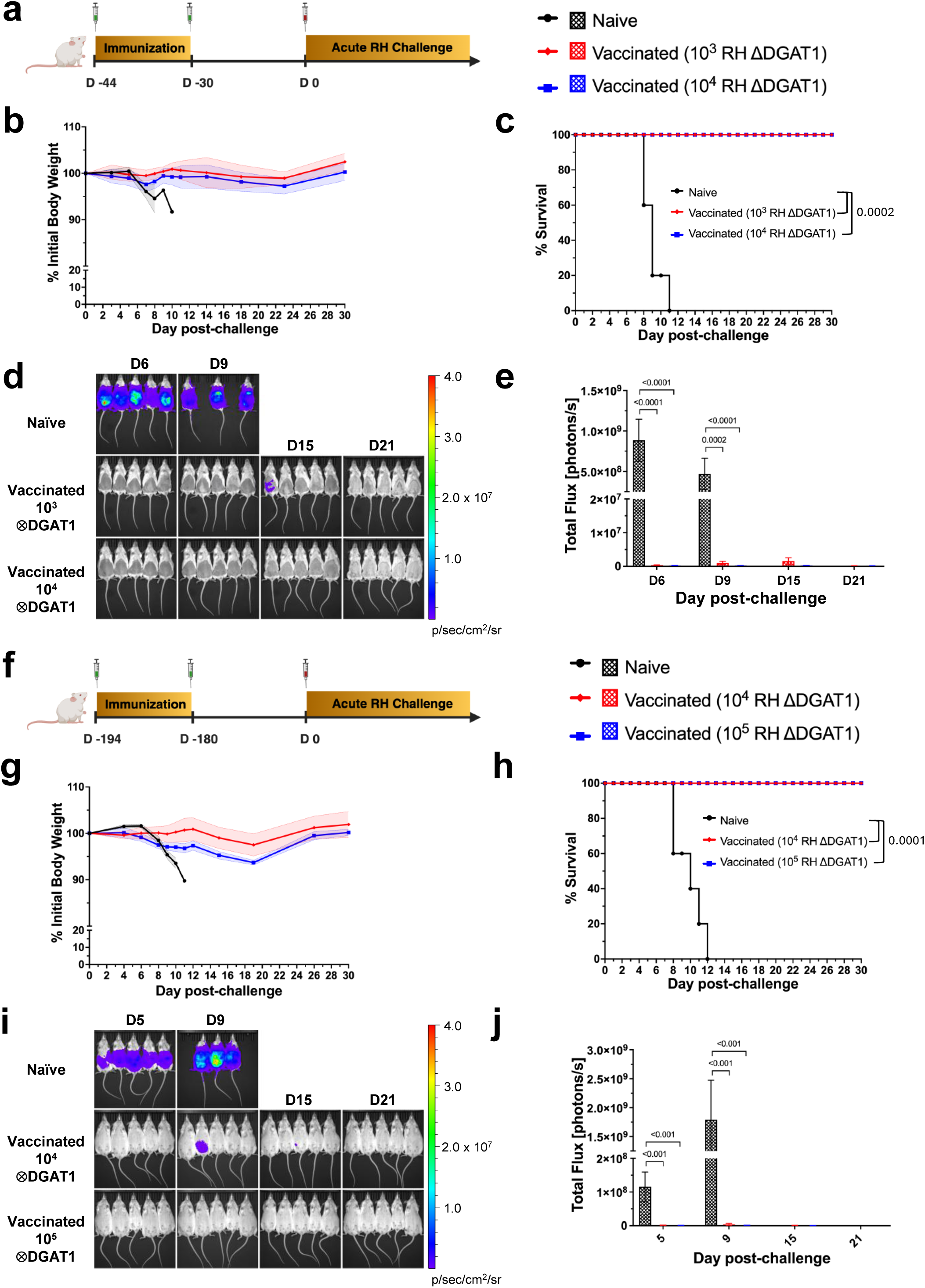
Vaccination with RH ΔDGAT1 parasites provides long-term protection of mice against acute type I *T. gondii* infection. (a) Female CD-1 mice (n = 5 per group) were given two immunization doses of 10^3^ or 10^4^ RH ΔDGAT1 parasites, spaced two weeks apart, followed by acute challenge with 500 tachyzoites of the type I RH luciferase-expressing *T*. *gondii* strain 30 days after the final immunization. (b) Normalized weights (means ± SEM) and (c) survival for naïve and immunized mice were plotted. Vaccinated mice showed no signs of infection or weight loss over 30 days, whereas naïve mice lost weight rapidly and died quickly. The log-rank (Mantel–Cox) test was used to assess the statistical significance of the difference in survival probability between naïve and vaccinated mice. (d) Bioluminescence images of naïve and immunized mice are shown at days 6, 9, 15, and 21 after infection with luciferase-expressing RH parasites. Vaccinated mice showed no parasite luminescence at day 21. (e) Quantification of whole-body luminescence of luciferase-expressing parasites in CD-1 mice, shown in (d). Mean ± SEM data (total flux, photons/sec) were plotted, and *p*-values were calculated using multiple unpaired t-tests with Holm-Sidak’s multiple-comparison test. (f) Female CD-1 mice (n = 5 per group) received two immunizations with 10^4^ or 10^5^ RH ΔDGAT1 parasites, spaced two weeks apart. They were then acutely challenged with 500 tachyzoites of the type I RH luciferase-expressing *T*. *gondii* strain 180 days after the last immunization. (g) Normalized weights (means ± SEM) and (h) survival for naïve and immunized mice were plotted. Vaccinated mice displayed no infection symptoms or weight loss during 30 days, while naïve mice quickly lost weight and died within 12 days of infection. The log-rank (Mantel–Cox) test was employed to evaluate the statistical significance of the survival probability difference between naïve and vaccinated mice. (i) Bioluminescence images of naïve and immunized mice are shown at days 5, 9, 15, and 21 post-infection with luciferase-expressing RH parasites. By day 21, vaccinated mice exhibited no detectable parasite luminescence. (j) Quantification of the total body luminescence from luciferase-expressing parasites in CD-1 mice, shown in (i). Mean ± SEM data (total flux, photons/sec) were plotted, and *p*-values were determined using multiple unpaired t-tests with Holm-Sidak’s correction for multiple comparisons.

We also tested the immunity conferred by RH ΔDGAT1 parasites against a heterologous type II *Toxoplasma* challenge (Figure 7a and f; Supplementary Figure 5a). We challenged inbred BALB/c mice, inoculated with 10^3^ to 10^5^ KO parasites, with a high lethal dose of 2000 luciferase-expressing Me49 tachyzoites (Me49-Luc). We tracked disease progression in infected mice over 30 days using bioluminescence imaging, while monitoring weight loss and mortality. The KO strain inoculation protected mice from the lethal Me49-Luc challenge at 30, 90, and 180 days. Uninoculated naive mice displayed clear clinical signs, rapid weight loss, high in vivo luminescence, and death within 12-13 days, while inoculated mice showed no obvious signs of acute *T*. *gondii* infection (Figure 7b-e, g-j; Supplementary Figure 5b). The cyst burden of the brains of all surviving mice was assessed at the time of euthanasia (30 days after challenge) based on rhodamine-labeled DBA staining. Due to the severe morbidity associated with the high Me49 challenge dose, we also harvested brains from all uninoculated moribund mice that survived more than two weeks after the challenge and reached their humane endpoint. No brain cysts were detected in mice challenged at 90 and 180 days post-inoculation (data not shown). In the 30-day challenge experiment, no cysts were observed in inoculated mice, except for one in the 10^3^ ΔDGAT1-inoculated group with just 160 cysts – likely due to faulty injection – while the euthanized naïve animals had an average of around 2,500 cysts (Supplementary Figure 5c). Dorsal IVIS imaging also revealed high luminescence in the brain region of naïve mice, while no detectable luminescence was found in the inoculated animals (Supplementary Figure 4d and e).

**Fig. 7.**
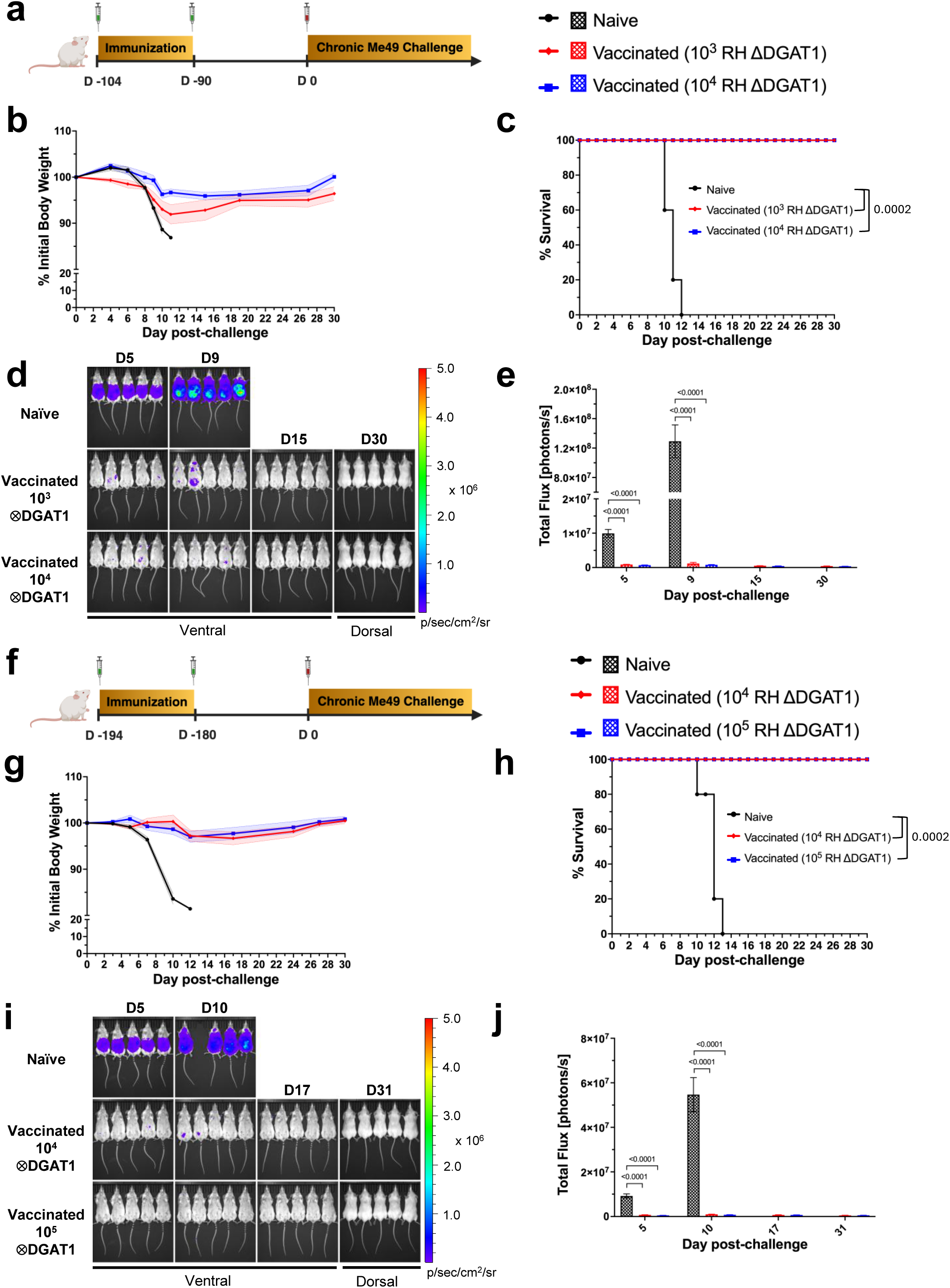
Immunization of mice with RH ΔDGAT1 parasites protects against cyst-forming type II *T*. *gondii*. (a) Female BALB/c mice (n = 5 per group) were given two immunization doses of 10^3^ or 10^4^ RH ΔDGAT1 parasites, spaced two weeks apart, followed by challenge with 2000 tachyzoites of the type II Me49 luciferase-expressing *T. gondii* strain 90 days after the final immunization. (b) Normalized weights (means ± SEM) and (c) survival rates for naïve and immunized mice are shown. Vaccinated mice did not exhibit signs of infection or weight loss for 30 days, while naïve mice lost weight rapidly and succumbed quickly. The log-rank (Mantel–Cox) test was used to assess the statistical significance of the difference in survival probability between naïve and vaccinated mice. (d) Bioluminescence images of naïve and immunized mice captured ventrally on days 5, 9, and 15, and dorsally on day 30 post-infection with luciferase-expressing Me49 parasites are shown. Vaccinated mice showed no parasite luminescence from day 15 onwards. (e) Measurement of the total body luminescence from luciferase-expressing parasites in BALB/c mice, shown in (d). Data are presented as mean ± SEM (total flux, photons/sec), and *p*-values were determined through multiple unpaired t-tests with Holm-Sidak’s correction for multiple comparisons. (f) Female BALB/c mice (n=5 per group) received two immunizations with 10^4^ or 10^5^ RH ΔDGAT1 parasites, administered two weeks apart, and were then challenged 180 days after the last dose with 2000 tachyzoites of the type II Me49 luciferase *T*. *gondii* strain. (g) Normalized weights (means ± SEM) and (h) survival rates for naïve and immunized mice are shown. Vaccinated mice showed no signs of infection or weight loss for 30 days, while naïve mice quickly lost weight and died. The log-rank (Mantel–Cox) test assessed survival differences. (d) Bioluminescence imaging of naïve and immunized mice on days 5, 10, 17 (ventrally), and 30 (dorsally) showed that vaccinated mice had no parasite luminescence from day 17. (e) Total body luminescence measurements in BALB/c mice, shown in (d), are presented as mean ± SEM (total flux, photons/sec). *p*-values were obtained using multiple unpaired t-tests with Holm-Sidak’s correction.

### Protection conferred by ΔTgDGAT1 immunization in mice is IFN-γ-dependent and predominantly CD8^+^ T cell-mediated

We next wanted to determine whether the resistance of mice to wild-type infection following inoculation with the mutant is due to a robust immune response mounted against the parasite, by examining which immune effectors could drive this protection. As a standard approach, we depleted essential immune components, such as IFN-γ, CD4^+^ T-cells, and CD8^+^ T-cells, in both naïve and inoculated mice using selective antibody-mediated depletion before challenging them with a type I infection (Figure 8a). Successful depletion of CD4^+^ T and CD8^+^ T-cells was confirmed before and after infection by flow cytometry (Supplementary Figure 6). As expected, the inoculated mice were shielded from infection, while naïve mice lost body weight and died within 9 days post-challenge (Figure 8b and c). Inoculated mice lacking IFN-γ also showed significant reductions in body weight accompanied by high mortality rates and ultimately succumbed to infection within 10 days after challenge. Furthermore, these mice had significantly higher infection levels (*P* = 0.0133) than naïve mice on day 7 after challenge, as measured by bioluminescence signal intensity (Figure 8d and e), indicating that immunity induced by the mutant inoculum depends heavily on IFN-γ. Depleting CD4^+^ T cells did not render inoculated mice susceptible to *T*. *gondii* RH strain infection. Interestingly, inoculated mice lacking CD8^+^ T cells exhibited delayed illness, resulting in the death of one mouse on day 15 and an 80% survival rate by day 30 post-challenge (Figure 8c). IVIS imaging showed peak infection levels in CD8^+^ T cell-depleted mice on day 11 post-inoculation, compared to the peak observed on day 7 in both naïve and inoculated mice depleted of IFN-γ (Figure 8c-e). Additionally, flow cytometry analysis of peripheral lymphocytes revealed an increase in CD8^+^ T cells in all inoculated mice on day 7 after *T*. *gondii* challenge, compared with pre-challenge levels (Supplementary Figure 6c). This indicates that CD8^+^ T cells, which are part of the adaptive immune system, play a crucial role in controlling *Toxoplasma* infection in RH ΔDGAT1-inoculated mice at later stages. Therefore, we refer to the RH ΔDGAT1-inoculated mice as RH ΔDGAT1-immunized or –vaccinated mice.

**Fig. 8.**
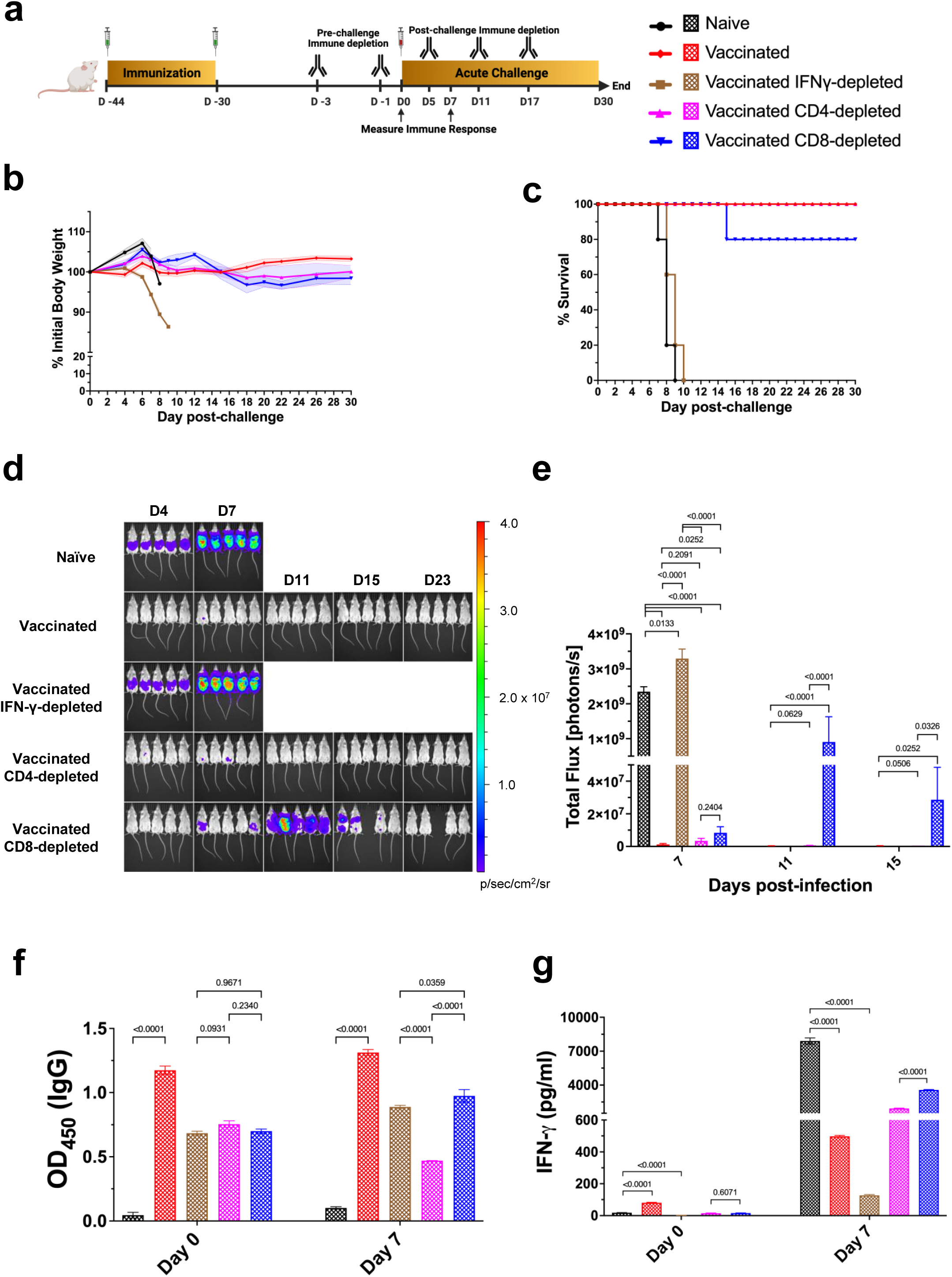
The immune response from ΔDGAT1 mutant vaccination in mice relies on IFN-γ and primarily engages CD8^+^ T cells. (a) Female BALB/c mice (n=5 per group) received two doses of 10^4^ RH ΔDGAT1 parasites, two weeks apart. IFN-γ levels, CD4^+^ T cells, or CD8^+^ T cells were then depleted via intraperitoneal injections of 200 μg of anti-IFN-γ (Clone XMG1.2), anti-CD4 (Clone GK 1.5), and anti-CD8b (Clone H35-17.2) monoclonal antibodies, or mock isotype controls (Clones HRPN IgG1 and LTF2 IgG2b) on specified days, before and after challenge with a lethal dose of 500 luciferase-expressing type I RH tachyzoites. IgG and IFN-γ responses were measured in the serum of naïve and vaccinated mice on day 0 and day 7 of challenge. (b) Normalized weights (means ± SEM) and (c) survival rates for naïve and immunized mice are shown. Vaccinated mice treated with isotype controls or anti-CD4 antibody showed no signs of infection or weight loss for 30 days, while naïve mice quickly lost weight and died by day 9 of infection. Mice depleted of IFN-γ also showed rapid weight loss and high mortality. Vaccinated mice depleted of CD8^+^ T cells showed delayed sickness, with one mouse dying on day 15. The log-rank (Mantel–Cox) test assessed survival differences. (d) Bioluminescence imaging of naïve and immunized mice at specific days showed no parasite luminescence in vaccinated mice treated with isotype controls or anti-CD4 antibody. High luminescence was observed in naïve and IFN-γ-depleted vaccinated mice, whereas vaccinated mice depleted of CD8^+^ T cells exhibited delayed infection. (e) Total body luminescence measurements in BALB/c mice, shown in (d), are presented as mean ± SEM (total flux, photons/sec). *p*-values were obtained using multiple unpaired t-tests with Holm-Sidak’s correction. (f) *Toxoplasma*-specific IgG levels were measured by ELISA on days 0 and 7 of the challenge. All vaccinated mice showed significantly higher IgG titers before the challenge than non-vaccinated mice. As anticipated, vaccinated mice depleted of CD8^+^ T cells exhibited the lowest IgG titers after the challenge. Mean ± SD data, analyzed by two-way ANOVA with Dunnett’s multiple-comparison test, are shown. (g) IFN-γ levels were measured by ELISA. As expected, IFN-γ levels were significantly reduced in IFN-γ-depleted vaccinated mice on both days of testing. On day 7 after the challenge, naïve mice showed the highest IFN-γ levels. Bars represent the means ± SD, with indicated *p*-values determined by multiple unpaired t-tests with Holm-Sidak’s multiple-comparison test.

We also measured *Toxoplasma*-specific total IgG and IFN-γ levels in sera collected from mice before and on day 7 of the challenge to assess the immune response in mice depleted of IFN-γ, CD4^+^ T cells, or CD8^+^ T cells. All immunized groups showed significantly higher pre-challenge IgG concentrations than the non-immunized group (*P* < 0.0001), regardless of immune depletion (Figure 8f). As anticipated, the immunized group exhibited the highest IgG concentrations both before and after the acute challenge. Depletion of IFN-γ, CD4^+^ T-cells, or CD8^+^ T-cells significantly reduced *Toxoplasma*-specific IgG concentrations (*P* < 0.0001). Although there were no statistical differences among IFN-γ-, CD4^+^ T-cell-, and CD8^+^ T-cell-depleted immunized mice before challenge, mice depleted of CD4^+^ T cells had significantly lower parasite-specific IgG levels (*P* < 0.0001) than the other groups on day 7 after the challenge (Figure 8f). This suggests that depletion of CD4^+^ T cells following RH ΔDGAT1 immunization hindered the development of *Toxoplasma*-specific antibodies, given their recognized role in sustaining antibody responses. However, it does not compromise the vaccine effectiveness to confer protective resistance against acute infection.

On the day of the challenge, IFN-γ levels were significantly elevated in vaccinated mice (*P* < 0.0001) compared to unvaccinated mice. Further, as expected, IFN-γ was undetectable in IFN-γ-depleted vaccinated mice (*P* < 0.0001) compared to unvaccinated mice, and this reduction persisted on day 7 after the challenge (Figure 8g). In all mice except those depleted of IFN-γ, post-challenge serum IFN-γ levels generally correlated with disease severity, with higher levels indicating more severe disease. We observed reduced IFN-γ in immunized mice compared to naïve mice and vaccinated mice depleted of CD4^+^ and CD8^+^ T cells. This indicates that RH ΔDGAT1 vaccination suppresses the severe host pro-inflammatory response during acute *Toxoplasma* infection. Managing host-specific immunity is vital for the host’s overall survival, as excessive immune responses can be harmful. Overall, vaccination with the RH ΔDGAT1 strain elicits an effective and safe protective immunity.

### B cells are required for ΔTgDGAT1 vaccine-induced immunity to acute infection

Studies using μMT mice lacking B cells revealed that B cells are essential for resisting chronic primary *T*. *gondii* infection and for conferring survival upon challenge with highly virulent parasites after vaccination (32, 33). To determine the role of antibodies or B cells in the resistance to *T*. *gondii* provided by RH ΔDGAT1 immunization, we infected BALB/c B cell-sufficient and B cell-deficient mice with RH ΔDGAT1 parasites and subsequently challenged them intraperitoneally after a month with highly virulent RH tachyzoites (Figure 9a). Both B cell-sufficient and B cell-deficient mice were highly vulnerable to acute type I *Toxoplasma* infection, with all animals losing body weight and succumbing within 8 days of challenge due to high parasite loads (Figure 9b and e). Immunized B cell-deficient mice, unlike immunized B cell-sufficient mice, were susceptible to RH challenge but experienced delayed infection and survived longer up to 22 days post-challenge (Figure 9b and e). Therefore, results suggest that inoculation helps B cell-deficient mice to combat parasite infection more effectively than naive mice; however, they ultimately died, unlike immunized mice having intact B cells. Parasite antigen-specific total IgG ELISA confirmed the absence of antibodies in the serum of naïve and immunized B cell-deficient mice before and after challenge, indicating that antibodies were necessary to block the infection in immunized animals (Figure 9f).

**Fig. 9.**
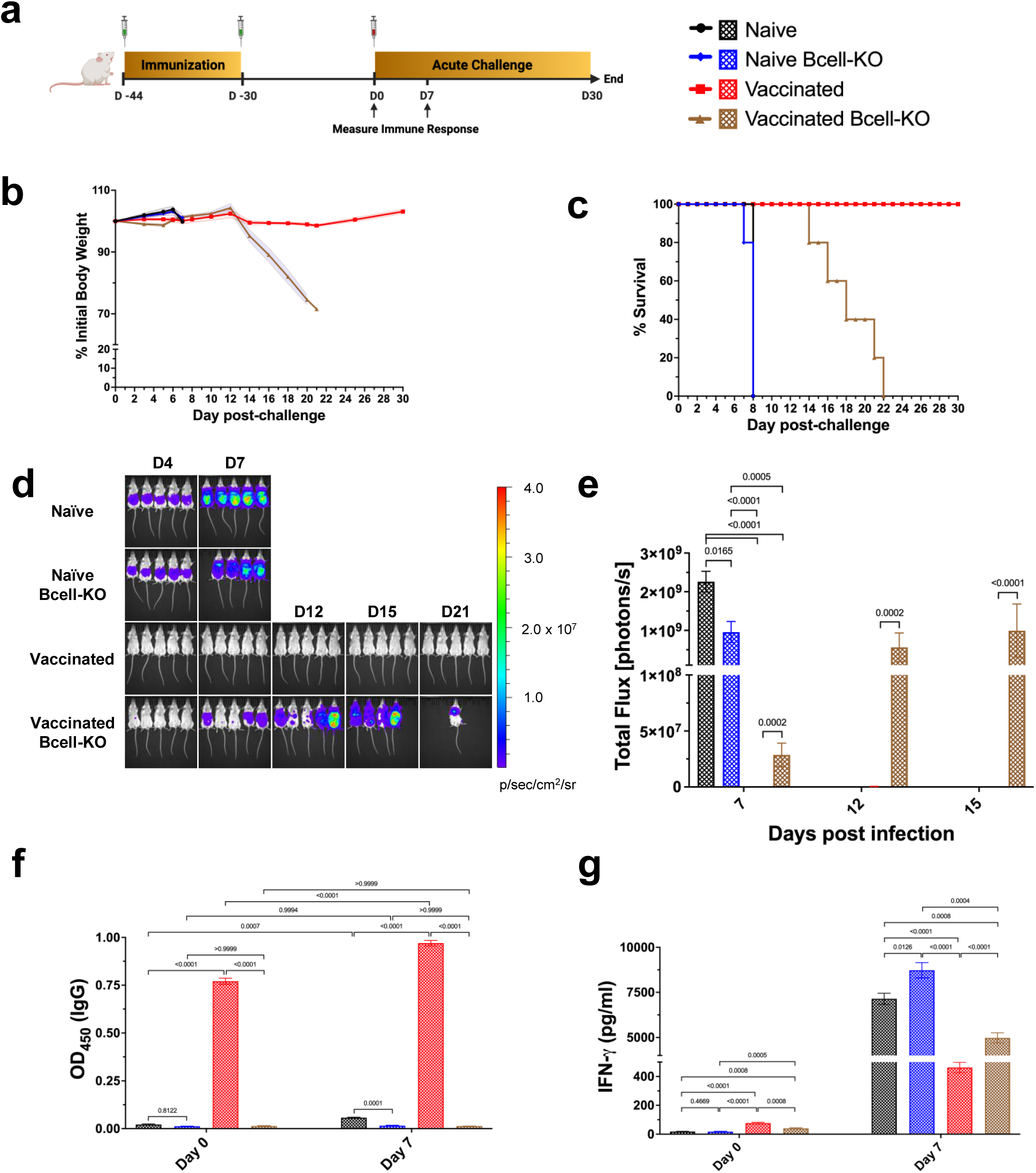
B cells are necessary for ΔDGAT1 vaccine immunity to acute infection. (a) Groups of five female BALB/c mice with or without a targeted deletion of B cells received two doses of 10^4^ RH ΔDGAT1 parasites, two weeks apart, followed by acute intraperitoneal challenge with 500 tachyzoites of luciferase-expressing type I RH *T*. *gondii* tachyzoites 30 days after the final immunization. IgG and IFN-γ responses were assessed in serum of naïve and vaccinated mice on days 0 and 7 of challenge. (b) Normalized weights (means ± SEM) and (c) survival for naïve and immunized mice were plotted. Both unvaccinated groups, whether B-cell-sufficient or B-cell-deficient mice, died within 8 days of infection. In contrast, vaccinated mice with B cells depleted showed slower disease progression and survived longer. The log-rank (Mantel–Cox) test assessed survival differences. (d) Bioluminescence images of naïve and immunized mice at specific days are shown. Both unvaccinated groups showed high parasite luminescence, while vaccinated B cell-sufficient mice had none. Vaccinated B-cell-deficient mice showed delayed infection. (e) Whole body luminescence measurements in BALB/c mice, displayed in (d), are shown as mean ± SEM (total flux, photons/sec). *p*-values were calculated using multiple unpaired t-tests with Holm-Sidak’s correction. (f) *Toxoplasma*-specific IgG levels were assessed by ELISA on days 0 and 7 of the challenge. Vaccinated mice with normal B cell function exhibited significantly higher IgG titers before and after the challenge compared to non-vaccinated mice. Mean ± SD data, analyzed by two-way ANOVA with Dunnett’s multiple-comparison test, are shown. (g) IFN-γ levels were assessed using ELISA. As anticipated, vaccinated mice maintained significantly lower IFN-γ levels after challenge, whereas naïve mice exhibited the highest levels. Data are shown as mean ± SD, with *p*-values obtained from multiple unpaired t-tests using Holm-Sidak’s correction.

Because IFN-γ is essential for resistance to acute RH tachyzoite challenge, we assessed IFN-γ production across all mouse groups. Naive and vaccinated mice were challenged with RH tachyzoites, and their sera were collected to measure IFN-γ levels before and after challenge by ELISA. On the day of challenge, IFN-γ level in vaccinated B-cell-deficient mice was significantly higher than those in unvaccinated B cell-sufficient and B cell-deficient mice (*P* = 0.0008 and 0.0005, respectively) but lower than in vaccinated B-cell-sufficient mice (*P* = 0.0008), suggesting that B cells might contribute to IFN-γ production in RH ΔDGAT1 vaccinated mice (Figure 9g). As previously observed in Figure 8, post-challenge serum levels of IFN-γ correlated with disease severity. Vaccination enabled B-cell-deficient mice to maintain lower pro-inflammatory IFN-γ levels compared to unvaccinated B cell-sufficient and B cell-deficient mice (*P* = 0.0008 and 0.0004, respectively), though these levels were not sufficiently reduced to prevent mortality as observed in B cell-sufficient mice (*P* < 0.0001). These findings suggest that the primary reason for the loss of resistance to *Toxoplasma* in vaccinated mice without B cells would be in part due to their inability to produce antibodies. However, secondary factors might include their reduced IFN-γ production prior to the challenge and challenges in maintaining lower IFN-γ levels afterward.

## Discussion

*Toxoplasma gondii* is auxotrophic for many essential metabolites, and its survival relies on its ability to acquire them from its host. In some instances, this parasite employs a highly effective dual-metabolic strategy for growth, combining de novo fatty acid synthesis and robust scavenging of host-derived fatty acids (34). These fatty acids are vital for membrane formation, energy production, and other metabolic functions. However, high levels of FFA can be toxic to the parasite. To avoid lipotoxicity, *Toxoplasma* produces lipid-esterifying enzymes that store excess FFA as neutral lipids, such as TAG and CE within intracellular LD. One such enzyme, TgDGAT1, catalyzes the final step of TAG synthesis via an acyl-CoA-dependent process. The DGAT1 inhibitor T863 inhibits *Toxoplasma* growth and replication in mammalian cells, indicating that TgDGAT1 plays a crucial role in storing energy-rich fatty acids as TAGs and safeguarding the parasite from FFA toxicity. We therefore studied the specific effects of DGAT1 loss in the parasite. Deleting DGAT1 caused defects in LD biogenesis, including significant replication loss, asynchronous replication, membrane defects, and a loss of virulence in both immunocompetent and immunocompromised mice, revealing the importance of this enzyme in *T*. *gondii*.

Previous research shows that vaccination can effectively control *Toxoplasma gondii*, with live attenuated vaccines demonstrating the most promise. Toxovax®, which utilizes the S48 “incomplete” strain of *T*. *gondii*, has been proven to reduce *Toxoplasma*-related abortions in sheep (11). Nevertheless, its application is mainly restricted to sheep due to concerns that the vaccine strain could revert to a virulent form. Currently, researchers are focusing on precise genetic modifications of parasites using the CRISPR/Cas9 system to delete critical parasite genes, eliminating the risk of reversion (35–50). Gene-edited live-attenuated vaccines are effective because they stimulate a broad immune response, similar to natural infections, providing long-lasting protection. In this study, we report that a *T*. *gondii* mutant deficient in a critical lipid storage enzyme (RH ΔDGAT1), when used as a whole-tachyzoite vaccine in inbred and outbred mice, provides cross-protection and long-term protection against subsequent lethal challenges by type I and type II *T*. *gondii* parasites.

Several type I *Toxoplasma*-based attenuated vaccines have previously shown effectiveness against acute lethal challenges of both type I and type II strains (36, 38, 40, 44, 46, 47, 50). However, these vaccines did not fully prevent cyst formation and chronic infection in mice challenged with a type II strain. In contrast, immunization with RH ΔDGAT1 parasites results in an undetectable cyst burden in challenged mice. It has also been proposed that using an attenuated type II vaccine strain could offer more reliable immunity against type II infection by blocking cyst development and chronic infection (35, 37, 39, 41, 42, 48, 49). However, these vaccines pose risks of reversion and cyst formation during vaccination, and they generally provide only partial protection against infection. Consequently, employing type I vaccine strains such as the RH ΔDGAT1, which offer full cross-protection against both type I and type II strains, might be the optimal strategy. The RH ΔDGAT1 mutant’s inability to cause disease, even in severely immunodeficient mice, suggests that parasites likely accumulate lipids over time in the body and die primarily from lipotoxicity rather than from direct host immune activity. Nonetheless, this persistence allows enough time for an immunocompetent individual to mount a strong response, ensuring protection against future infections even after extended periods post-vaccination. Because the type I ΔDGAT1 RH strain is slow-growing, entirely avirulent, and non-cystogenic, the immune response triggered by vaccination with this strain might explain its improved control of type II chronic infection. However, the specific mechanism underlying this cross-protection requires further detailed investigation.

We examined both innate and adaptive immune responses, including humoral and cellular responses, associated with protection against *Toxoplasma* infection following parenteral immunization with the genetically attenuated RH ΔDGAT1 vaccine. Our data suggest that immunity against *T*. *gondii* from RH ΔDGAT1 vaccination results from both cellular and humoral immune responses. The serum *Toxoplasma*-specific antibody levels observed in this study were relatively higher and persisted for longer than those typically reported for other live attenuated vaccine platforms. In this study, high ELISA OD values indicating elevated IgG concentrations were detected even at an extremely high serum dilution of 1:24,300 whereas other studies typically use serum dilutions of 1:50 to 1:100. We evaluated the production of Th1 (proinflammatory) cytokines IFN-γ, IL-12, TNF-α, and Th2 (anti-inflammatory) cytokines IL-4 and IL-10 in the splenocytes of the mice after in vitro recall with antigens (STAg). The levels of both cytokine types stayed significantly higher at 180 days after vaccination compared to the control, showing that the ΔDGAT1 vaccine induces a regulated immune response. While inflammation helps control parasites, overly strong, uncontrolled immune responses can damage tissue through a lethal inflammatory process, underscoring the need for a balanced response that eliminates parasites while minimizing collateral injuries (51). Interestingly, our findings also indicate that the dose of the ΔDGAT1 vaccine primarily shapes the Th1/Th2 response profile. Specifically, early in vaccination, lower doses mainly induce a cell-mediated Th1 response, while higher doses lead to a mixed Th1/Th2 response. A similar pattern was observed previously with the Bacille Calmette-Guérin (BCG) vaccine against *Mycobacterium tuberculosis*, which is also an intracellular vacuolar pathogen like *T*. *gondii* (52).

Multiple studies clearly show that CD8^+^ T cells and IFN-γ are vital for defending against *Toxoplasma* infection in mice (25–28, 53, 54). Our immune depletion experiments in mice supported this finding. Additionally, we observed that RH ΔDGAT1-vaccinated mice maintained high levels of IFN-γ in their splenocytes for at least 6 months post-vaccination. Effector memory CD44^+^CD62L^-^ T cells were more prevalent and persisted longer in ΔDGAT1-vaccinated mice, demonstrating their superior ability to manage infection during the challenge even 180 days post-vaccination.

It has been shown that mice lacking CD4^+^ T cells, which cannot produce isotype-switched antibodies, exhibit reduced protection against challenge with virulent RH tachyzoites after being immunized with the attenuated ts-4 strain of *T*. *gondii* (55). This contrasts with our results, which suggest that CD4^+^ T cells are dispensable for vaccine protection. Notably, those researchers immunized CD4-depleted mice, whereas we immunized wild-type mice and then depleted CD4^+^ T cells before challenge. This suggests that CD4^+^ T cells may be crucial only during vaccination and not after anti-*Toxoplasma* immunity is established. In fact, a study reported that anti-CD4 antibodies can hinder protective immunity when administered during vaccination rather than after challenge (53).

Previous research indicates that B cell-deficient mice exhibit reduced resistance to *Toxoplasma gondii* infection, despite unaffected expression of IFN-γ, TNF-α, and inducible nitric oxide synthase (33). Further, in addition to cell-mediated immunity, B cells are crucial for protection against an intraperitoneal challenge with virulent *Toxoplasma* parasites in mice vaccinated with attenuated tachyzoites (32). This protection is humoral in nature, as the passive transfer of antibodies confers protection to B cell-deficient mice (55). Moreover, resistance to secondary *T*. *gondii* infection is linked to a robust, regulated humoral response in resistant mice, compared with that in highly susceptible mice (56). Our results also indicate that B cells are crucial for the antibody-mediated immunity conferred by the ΔTgDGAT1 vaccine against acute RH infection, with a possible minor role in IFN-γ production. However, B cell-deficient mice vaccinated with ΔTgDGAT1 and challenged with RH tachyzoites lived noticeably longer than unvaccinated mice. This implies that even without B cells, some resistance – likely mediated by T cells and dependent on IFN-γ – is still generated, though it is insufficient for full protection. Nonetheless, vaccination can help resist a more physiologically relevant challenge infection with a mildly virulent type II strain of *T*. *gondii* in the absence of B cells (57). Whether ΔTgDGAT1 vaccination can protect B-cell-deficient mice against a type II infection remains to be studied.

Our work using the CRISPR-Cas9 system demonstrates that deleting the *DGAT1* gene from the highly virulent *Toxoplasma* type-I RH strain results in lipid droplet depletion, thereby reducing the parasite’s ability to survive inside host cells. Notably, even at lethal doses, the RH ΔDGAT1 strain remains non-virulent in IFN-γ-KO and triple-immunodeficient NCG mice, showing significant attenuation of virulence. Validating TgDGAT1 as a drug target in our lab has inspired us to find specific inhibitors for this crucial lipid-esterifying MBOAT that block parasite growth. The observation that existing lipid storage inhibitors can affect *Toxoplasma* growth is also intriguing and merits further research.

Further studies are needed to assess the safety and effectiveness of the ΔDGAT1 mutant in food animals such as sheep and in the primary feline host. Our assumption that this vaccine holds the premise to interrupt the parasite’s life cycle at its source by reducing infective cysts and oocysts, thereby minimizing economic losses in livestock, protecting the human food supply, and preventing severe congenital diseases in pregnant women. Indeed, infection with *T*. *gondii* before pregnancy can induce protective immunity, highlighting the advantages of vaccinating before conception (58).

Overall, our results offer important insights into how the highly effective RH ΔDGAT1 vaccine induces immune protection. This information can be compared with data from other *Toxoplasma* vaccines to guide the development of *Toxoplasma* vaccines and, potentially, those for other key apicomplexan parasites. RH ΔDGAT1 emerges as a promising candidate for a live-attenuated vaccine to combat toxoplasmosis.

## Methods

### Ethics

All procedures were conducted in accordance with the Public Health Service Policy on Humane Care and Use of Laboratory Animals and the Association for the Assessment and Accreditation of Laboratory Animal Care guidelines. The animal protocol involving infecting mice with *T*. *gondii* was approved by the Institutional Animal Care and Use Committee at Johns Hopkins University School of Public Health.

### Reagents and antibodies

All common chemicals and reagents were obtained from Sigma-Aldrich (St. Louis, MO) unless otherwise stated. [5,6-^3^H] uracil (ART 0282) was purchased from American Radiolabeled Chemicals, Washington, D.C. The mouse monoclonal anti-SAG1 primary antibody used for immunofluorescence was a generous gift from J.-F. Dubremetz, Université de Montpellier. The goat anti-mouse secondary antibody conjugated to Alexa Fluor 488 (A11029) was purchased from Invitrogen (Waltham, MA).

### Mammalian cells, parasites and culture conditions

Human foreskin fibroblasts (HFF-1, SCRC-1041) were obtained from the American Type Culture Collection (Manassas, VA). All cells were grown in α-minimum essential medium (α-MEM) supplemented with 10% fetal bovine serum (FBS), 2 mM L-glutamine, 100 U/mL penicillin, and 100 μg/mL streptomycin unless stated otherwise, and maintained at 37°C with 5% CO_2_. *Toxoplasma gondii* tachyzoites used in this study included the type I (RH) and the type II (Me49) strains. The RH Δku80Δhxgprt (ATCC PRA-319) (17) (WT) strain was obtained from V. Carruthers (University of Michigan), while the luciferase-expressing RH Δhxgprt-GFPLuc (59) and Me49 Δhxgprt-Fluc (60) strains were kindly provided by M. Grigg (National Institute of Allergy and Infectious Diseases) and L. Knoll (University of Wisconsin-Madison), respectively. These parasites were propagated in vitro by serial passage in monolayers of HFF-1 cells grown in α-MEM supplemented with 10% FBS, 2 mM L-glutamine, 100 U/mL penicillin, and 100 μg/mL streptomycin. The *T*. *gondii* RH ΔDGAT1 (KO) strain created in this study was maintained in HFF-1 using α-MEM supplemented with 10% lipid-depleted FBS (LPD-FBS, Omega Scientific), 3.5 g/L glucose, 2 mM L-glutamine, and penicillin/streptomycin at 37°C and 5% CO_2_. For preparation of OA-containing medium, sodium oleate was dissolved in sterile water at a concentration of 100 mM at 50°C and then thoroughly mixed by vortexing with prewarmed 5% fatty acid-free bovine serum albumin (BSA) in a 1:15 ratio to ensure OA-BSA complex formation. The OA:BSA mixture was then added to prewarmed α-MEM to prepare growth media containing 0.4 mM OA and 0.3% BSA (OA-MEM) with an approximate 6:1 OA:BSA molar ratio.

### Mice

All mice in this study were female and kept in the animal care facility at Johns Hopkins Bloomberg School of Public Health. CD-1 IGS (Crl:CD1(ICR)), BALB/c (BALB/cAnNCrl), and NCG (NOD-Prkdc^em26Cd52^Il2rg^em26Cd22^/NjuCrl) mice were purchased from Charles Laboratories (Wilmington, MA) and used at 6 to 8 weeks of age. IFN-γ KO mice (B6.129S7-Ifngtm1Ts/J), aged 7 weeks, were procured from The Jackson Laboratory (Bar Harbor, ME). B cell-deficient mice (C.Cg-*Igh-J*^tm1Dh^u) in the BALB/c background (Jh) were purchased from Taconic Biosciences (Rensselaer, NY). Mice were acclimated to the local environment for at least one week before the experiments. Unless otherwise stated, five mice were used per group.

### Creation of the mutated strains by CRISPR/Cas9

The CRISPR/Cas9 gene editing system was used to delete the *Tg*DGAT1 gene in the RH Δku80Δhxgprt strain background. The plasmid pTOXO_Cas9-CRISPR::sgDGAT1 vector was generated as previously described (61). Briefly, the sense and anti-sense oligos sgRNATgDGATFw (5′-AAGTT**ATGTCGGTTGTGGAATCGAAG**G-3′) and sgRNATgDGATRev (5′-AAAAC**CTTCGATTCCACAACCGACAT**A-3′) containing the sgRNA (highlighted in bold) targeting the *Tg*DGAT1 genomic sequence (TGGT1_232730) were annealed, phosphorylated, and ligated in the pTOXO_Cas9-CRISPR plasmid linearized with BsaI, yielding pTOXO_Cas9-CRISPR::sgDGAT1, which was confirmed by DNA sequencing. To generate the Δ*Tg*DGAT1 targeting construct for homologous recombination repair, we amplified the selection cassette (LoxP-5′dhfr/HXGPRT (minigene)/3′dhfr-LoxP) from the pTKO vector (gift of M. Grigg, NIH) using primers P5 (5′-**CAGGGCGTATCGATTCTAGCTTTGACAGACTGTCCTAAAT**TTTGTACAAAAAAGCAGGCT-3′) and P6 (5′-**CTACATGATCTGTATTTTGCTCGGGTCTAGCTGCTGTACC**CCACAGTGAGTATCTCCCAC-3′) that contained 40 bp sequences (highlighted in bold) homologous to the genomic 5′ UTR sequence and the genomic sequence from the last exon of TGGT1_232730. The PCR product was purified and sequenced to confirm its identity. Subsequently, RH Δku80Δhxgprt parasites (1 × 10^7^) were mixed with 8.5 μg of pTOXO_Cas9-CRISPR::sgDGAT1 and 30 μg Δ*Tg*DGAT1 targeting construct (PCR product) and electroporated in cytomix buffer (120 mM KCl, 0.15 mM CaCl_2_, 10 mM K_2_HPO_4_/KH_2_PO_4_, 25 mM HEPES, 2 mM EDTA, 5 mM MgCl_2_, pH 7.6) as described previously (62). Selection of transfected parasites was initiated 24 h post-transfection using mycophenolic acid (25 μg/ml) and xanthine (50 μg/ml). Transfected parasites were single-cell cloned by limiting dilution in 96-well plates. Clonal lines were genotyped by PCR analysis. Genomic DNA was extracted from parasite clones using the QuickExtract DNA Extraction Solution (QE09050, Lucigen) as per the manufacturer’s instructions. PCR was performed on the extracted DNA to confirm the intended deletion of *Tg*DGAT1 after homology-directed repair as described in Supplementary Figure 1. Primers for screening the deletion included one set internal to the sequence to be deleted (P1; 5′-TCTAGGTGCATTCGTAATCGAG-3′ and P2; 5′-GTTCTGTGGTCCTCACGTAC-3′), another upstream and downstream of the gene deletion (P3; 5′-GAGGACGACTTGACCTGAAG-3′ and P4; 5′-AGTGTCAAGGTCAGTGACAC-3′), and the primer pair (P5/P6) used to generate the targeting construct.

### Parasite replication assays

Tachyzoite replication was assayed by either uracil incorporation assays or parasite enumeration per PV. For tritiated uracil incorporation assays, HFF-1 cells were grown to confluence in 24-well plates before infection with 1 × 10^5^ parasites for 4 h at 37 °C. Cells were washed twice with PBS and incubated for 24 h in αMEM medium. Cells were then incubated with 1 μCi of [5,6-^3^H] uracil for 2 h at 37 °C, and the samples were processed as described previously (63). To assess parasite number per PV, coverslips with confluent HFF-1 were infected with parasites for 4 h and thoroughly washed with PBS to remove extracellular parasites. Coverslips were fixed and stained as described above and viewed with a Zeiss AxioImager M2 fluorescence microscope equipped with an oil-immersion Zeiss plan APO 100×/NA 1.4 objective. The number of parasites was recorded for at least 100 PVs on the coverslip for each condition and the mean percentage PV distribution was recorded.

### Parasite growth assays

To monitor tachyzoite growth and development, plaque assays were performed using HFF-1 grown until confluence in 6-well plates. Each well was infected with *Toxoplasma* RH parasites, and plates were incubated at 37°C for 10 or 14 days without disturbance. The cells were fixed with 100% ethanol for 5 min at room temperature, stained with crystal violet stain solution (1% ammonium oxalate, 10% crystal violet in 100% ethanol) for 5 min and washed 4 times with PBS. The plates were scanned using a ScanWizard 5 scanner (Microtek) or imaged using Cytation 7 cell imaging system (Agilent), and the number and area of each plaque were measured using Fiji ImageJ with the freehand tool.

### Immunofluorescence assays and fluorescence microscopy

For immunolabeling, infected HFF-1 cells were fixed in PBS with 4% formaldehyde (Polysciences, Warrington, PA) and 0.02% glutaraldehyde for 15 min, permeabilized with 0.3% Triton X-100 in PBS for 5 min, and washed twice with PBS before blocking. Cells were incubated in blocking buffer (3% BSA, fraction V; Thermo Fisher Scientific, in PBS) for 1 h, followed by incubation with the anti-SAG1 primary antibody diluted (1:500) in blocking buffer for 1 h or overnight. Cells were washed three times with PBS for 5 min each and incubated in secondary antibody diluted (1:2,000) in blocking buffer for 1 h, followed by three washes with PBS for 5 min each. Cells were then incubated with a 1:1,000 dilution of 1 mg/ml DAPI (Roche) in PBS for 5 min, followed by 3 washes with PBS. To stain neutral lipids, cells were also incubated with a 1:1,000 dilution of LipidTOX Red (Invitrogen, Waltham, MA) in PBS for 30 minutes, then washed three times with PBS. Coverslips were rinsed with water and mounted on slides with ProLong Glass Antifade mounting solution (Invitrogen, Waltham, MA). Fixed samples were viewed with a Zeiss AxioImager M2 fluorescence microscope equipped with an oil-immersion Zeiss plan APO 100×/NA 1.4 objective, a Hamamatsu ORCA-R2 camera and Volocity software (Quorum Technologies, Ontario, Canada) or a Leica DMi8 Thunder microscope equipped with an oil-immersion HC PL APO 63×/NA 1.4 objective or HC PL APO 100×/NA 1.4 objective, a Leica K8 Scientific sCMOS camera and LAS X software.

### Fluorescence image analysis

Images were deconvolved with an iterative restoration algorithm using calculated point-spread functions, a confidence limit of 100% and an iteration limit of 30–35 using Volocity software. Images were cropped and adjusted for brightness and contrast using Volocity software.

To assess the number of LD associated with the parasites, optical z-sections of infected cells were acquired, deconvolved, and measured using the following measurement protocols in Volocity: the parasites were identified with the ‘Find Objects Measurement’ tool using *Tg*SAG1 fluorescence; thresholds were set manually using standard deviation (SD) with a lower limit of 0 and a minimum object size of 5 μm^3^. To enclose the identified SAG1 staining as a complete parasite volume, the object was processed with 30 iterations of ‘Close’ and ‘Fill Holes in Objects’. LD were identified with the ‘Find Objects Measurement’ tool based on LipidTox Red fluorescence with thresholding using SD intensity (lower limit of 2.5) and a minimum object size of 0.05 μm^3^. The SD threshold was set to begin at the center of the binomial curve to increase the inclusion of smaller LD (by size and intensity), especially the ones within the parasites. Noise was removed using a fine filter, and objects were separated using the ‘Separate Touching Objects’ tool in Volocity with an object size guide of 0.5 μm^3^. Following the pre-defined criteria, objects of size < 0.05 µm^3^ were excluded through ‘Exclude Objects by Size’. After performing a manual analysis of the detected objects, a ‘Filter Population’ was applied to enhance the population with the LD and eliminate non-LD specific staining. Lastly, the ‘Compartmentalize’ tool was used to determine the number of LD inside the parasites.

### Parasite ultrastructural observations

For ultrastructural observations by thin-section transmission electron microscopy, infected cell monolayers were washed 3 times with PBS, fixed in 2.5% glutaraldehyde (Electron Microscopy Sciences, Hatfield, PA) in 0.1 mM sodium cacodylate (pH 7.4) for 1 h at room temperature and processed as described (64) before examination with a Hitachi 7600 electron microscope under 80 kV, equipped with a dual AMT CCD camera system.

### Murine virulence assays

To evaluate the role of *Tg*DGAT1 in *T*. *gondii* virulence, groups of BALB/c, IFN-γ KO, and NCG mice were infected with 250 tachyzoites of WT or KO strains intraperitoneally (i.p.) in a volume of 200 μL sterile PBS. Clinical signs and mortality of infected mice were monitored daily for 30 days. Mice were also weighed at regular intervals till study completion.

### Quantitative PCR for estimation of parasite burden in mice

To assess parasite burdens in infected mice, groups of BALB/c mice (n=3 per group) received i.p. injections of 250 WT or KO tachyzoites. After 7 days, the mice were euthanized, and samples of peritoneal lavage fluid (1 ml), liver (20 mg), and spleen (10 mg) tissues were collected. Additionally, KO parasite loads in these tissues were measured in both BALB/c and NCG mice (n=3 per group) 30 days post-infection. The genomic DNA from the peritoneal fluid, liver, and spleen was extracted using the GenElute Mammalian Genomic DNA Miniprep Kit (Sigma-Aldrich, St. Louis, MO) and subjected to qPCR with primers targeting the *Toxoplasma* 529-bp repeat region (65, 66) (Tox-9F: 5′-AGGAGAGATATCAGGACTGTAG-3′; Tox-11R: 5′-GCGTCGTCTCGTCTAGATCG-3′) and the murine β-actin gene (Mouse Actin-F: 5′-GGCTGTATTCCCCTCCATCG-3′; Mouse Actin-R: 5′-CCAGTTGGTAACAATGCCATGT-3′). The qPCR was performed using the PowerUp SYBR Green Master Mix (Applied Biosystems, Waltham, MA) with 500 nM of each primer and 40 ng of DNA in a QuantStudio 6 Pro Real-Time PCR System (Applied Biosystems, Waltham, MA). Parasite burdens were calculated based on a standard curve generated from ten-fold serial dilutions of tachyzoite genomic DNA (4 × 10^7^, 4 × 10^6^, 4 × 10^5^, 4 × 10^4^, 4 × 10^3^, 400, and 40 tachyzoites), and corresponding cycle threshold (CT) values of the 529-bp repeat element normalized to the mouse β-actin gene.

### Measurement of *Toxoplasma*-specific IgG and cytokine levels

BALB/c mice were inoculated i.p., two weeks apart, with two doses of freshly harvested 10^4^ or 10^5^ KO tachyzoites in 200 μl PBS or mock-vaccinated with PBS. Serum samples were collected by retroorbital bleed to measure *Toxoplasma*-specific IgG and cytokines by ELISA at 30, 90, and 180 days after vaccination in vaccinated and naïve mice.

For total *Toxoplasma*-specific IgG and IgG subclass detection, RH WT tachyzoites were harvested, purified by filtration, and resuspended in ice-cold PBS. The resuspended parasites were lysed by sonication on ice, and the cleared soluble fraction of the lysate containing *T*. *gondii* antigens was clarified by centrifugation at 12,000 × g for 15 minutes at 4 °C. 96-well immuno plates were coated with soluble parasite antigens (1 μg per well), diluted in ELISA coating buffer (BioLegend, San Diego, CA), and incubated overnight at 4°C. The antigen-coated ELISA plates were washed twice with 0.05% Tween-20 in PBS (PBS-T), then once with PBS. Non-specific binding sites were blocked with 1% BSA (in PBS) for 1 hour and washed again. Serum samples were diluted in 1% BSA at a 1:300 ratio, and threefold serial dilutions were prepared in 1% BSA, with dilutions of 1/900, 1/2700, 1/8100, 1/24300, and 1/72900. Diluted serum samples were then added to wells and incubated for 2 h. The plates were then washed and HRP-conjugated goat anti-mouse IgG, IgG1, and IgG2a (Jackson ImmunoResearch, West Grove, PA) secondary antibodies (1:5000 dilution in 1% BSA) were added and incubated for 1 h in the dark. After washing, TMB High Sensitivity Substrate Solution (BioLegend, San Diego, CA) was added to wells for 10 min for color development, followed by the addition of Stop Solution for TMB Substrate (BioLegend, San Diego, CA), and measurement of optical density (OD) at 450 nm using a microplate reader. The expression levels of cytokines interferon-gamma (IFN-γ), interleukin 12p70 (IL-12p70), tumor necrosis factor alpha (TNF-α), interleukin 4 (IL-4), and interleukin 10 (IL-10) were detected in splenocytes isolated from vaccinated and naïve mice sacrificed at 180 days after vaccination. Spleens were collected aseptically from mice euthanized by CO_2_ overdose and cervical dislocation. The harvested spleens were minced into small fragments (∼0.2 cm²) using a razor, then incubated for 30 minutes at 37°C in RPMI-1640 medium supplemented with Collagenase IV (100 U/mL), DNase I (20 U/mL), and 10% FBS. Splenocytes were passed through a 70μm cell strainer, washed with PBS, hemolyzed in ice-cold 1× RBC Lysis Buffer (Invitrogen, Waltham, MA) for 5 minutes, and washed again with PBS to obtain a single spleen cell suspension. Subsequently, splenocytes were resuspended in RPMI-1640 culture medium with 10% FBS, and their viability and count were determined using trypan blue staining. Then, 2 × 10^6^ viable cells were cultured in RPMI-1640 medium supplemented with 10% FBS, 100 U/mL penicillin, and 100 μg/mL streptomycin in 24-well plates and stimulated with 10 μg/mL soluble tachyzoite antigens from the RH WT strain. Cell-free supernatants were collected at the specified time points—24 hours for IL-4, 72 hours for TNF-α and IL-10, and 96 hours for IL-12p70 and IFN-γ—and then stored at –80°C for cytokine analysis. Cytokines were detected in diluted or undiluted supernatants by using ELISA MAX Deluxe kits (BioLegend, San Diego, CA) according to the manufacturer’s recommended protocols.

### Type I and type II *Toxoplasma* challenge infections

CD-1 or BALB/c mice were vaccinated twice by i.p. injection, two weeks apart, with 10^3^, 10^4^, or 10^5^ KO parasites, as indicated, in 200 µl PBS, or mock-vaccinated with the same volume of PBS. Naïve and immunized CD-1 mice were challenged at 30, 90, or 180 days post-immunization with a lethal i.p. dose of 500 luciferase-expressing type I RH tachyzoites to assess protection against acute fatal infection over both short– and long-term periods. The disease progression was monitored in infected mice over 30 days by bioluminescence imaging and tracking weight loss and mortality. For in vivo bioluminescence imaging, mice received an i.p. injection of 200 μL of 15 mg/mL luciferin (D-Luciferin potassium salt, MediLumine, Quebec, Canada), then were immediately anesthetized in an oxygen-rich induction chamber with 2.5% isoflurane. Whole-body luminescence was measured in ventral position ten minutes after the luciferin injection using a Xenogen IVIS Spectrum Imaging System (Caliper Life Sciences, Hopkinton, MA) and analyzed with the Living Image software.

To assess whether the vaccine could prevent cyst formation following a type II *Toxoplasma* challenge, both vaccinated and unvaccinated BALB/c mice were exposed to a lethal dose of 2000 luciferase-expressing Me49 parasites at 30, 90, or 180 days post-vaccination. Over 30 days, challenged mice were monitored via in vivo imaging, weight loss, and recording survival. Mice were injected intraperitoneally with 200 μL of 1 mg/mL CycLuc1 (MedChemExpress, Monmouth Junction, NJ), and anesthetized mice were imaged in the ventral position as described above. For brain imaging, mice were positioned dorsal side up, and imaging was performed 20 minutes after injection of CycLuc1. Additionally, brains were harvested from surviving mice (30 days after challenge) or severely moribund mice (starting 2 weeks after challenge) to assess the cyst load using Rhodamine-labeled Dolichos biflorus agglutinin (DBA) staining. Whole brains were homogenized in 2 mL ice-cold PBS by repeatedly passing them through 16-gauge, 18-gauge, and 21-gauge needles until the mixture was uniform. The final homogenate volume was recorded, and one-tenth of this volume was combined with 800 µL of cold 100% methanol, then fixed at room temperature for 5 minutes. The samples were centrifuged at 5,000 × g for 5 minutes, washed with 1 mL of PBS, centrifuged again, and the supernatant was aspirated. To label cysts, the pelleted samples were incubated in 1 mL of PBS containing 10 µg/mL Rhodamine-conjugated DBA (Vector Laboratories, Newark, CA) with rotation at room temperature using a HulaMixer® Sample Mixer (Invitrogen, Waltham, MA). After 1 hour, the samples were washed twice with PBS, resuspended in 1 mL of PBS, and four 50-µL aliquots were plated into a clear-bottomed 96-well plate. These were examined under an inverted fluorescence microscope with a 10× objective to count DBA-positive tissue cysts. Brain cyst burdens were calculated as the average of the four counts per sample multiplied by 200.

### Cell depletions and flow cytometry

For in vivo depletion of IFN-γ, CD4^+^ T cells, and CD8^+^ T cells in RH Δ*Tg*DGAT1-vaccinated mice, 200 μg of rat anti-IFN-γ (Clone XMG1.2, BioXCell, Lebanon, NH), rat anti-CD4 (Clone GK 1.5, BioXCell, Lebanon, NH), rat anti-CD8b (Clone H35-17.2, Invitrogen, Waltham, MA) monoclonal antibodies, or rat isotypes controls (Clones HRPN IgG1 and LTF2 IgG2b, BioXCell, Lebanon, NH) were injected intraperitoneally on the indicated days before and after challenge with a lethal dose of 500 luciferase-expressing type I RH tachyzoites to identify the mediators of vaccine immunity against *Toxoplasma* infection. To assess the proportions of effector memory CD4^+^ and CD8^+^ T lymphocytes, peripheral blood samples were collected via retro-orbital bleeding. Cells were separated by centrifugation, and red blood cells were lysed with ACK Lysing Buffer (Quality Biological, Gaithersburg, MD). Isolated lymphocytes were kept on ice and stained immediately with a panel of antibodies for 30 minutes in the dark, targeting the following surface molecules: AF700-labeled anti-mouse CD3 (Clone 17A2, BioLegend, San Diego, CA), PE-labeled anti-mouse CD4 (Clone H129.19, BD Pharmingen, San Diego, CA), APC-labeled anti-mouse CD8a (Clone 53-6.7, Invitrogen, Waltham, MA), PE-Cy7-labeled CD44 (Clone QA19A43, BioLegend, San Diego, CA), and APC-Cy7-labeled CD62L (Clone MEL-14, BioLegend, San Diego, CA). The cell suspension was washed twice with PBS containing 2% FBS, then fixed with PBS containing 2% formaldehyde. All samples were analyzed by flow cytometry using the Attune Nxt Flow Cytometer (Invitrogen, Waltham, MA), and data were processed with FlowJo software Version 11 (Tree Star Inc., Ashland, OR). Lymphocyte populations were identified from blood mononuclear cells using forward-scatter (FSC) and side-scatter (SSC) parameters. Subsequently, the T cell population was isolated from these lymphocytes based on their CD3 surface expression. We identified CD4^+^ and CD8^+^ T cells using flow cytometric phenotypes, and memory effector T cells were identified by a CD44^+^CD62L^−^ phenotype.

### Statistical analyses

All statistical analyses were conducted with GraphPad PRISM v11. Data normality was evaluated using Q-Q plots and the Shapiro-Wilk test. Statistical comparisons between the groups were conducted using unpaired two-tailed t-tests, one-way ANOVA, or two-way ANOVA, with post hoc tests such as Holm-Sidak’s, Tukey’s, or Dunnett’s, depending on the data. The log-rank (Mantel–Cox) test was used to assess the statistical significance of the difference in survival probability between naïve and vaccinated mice. *P* values of 0.05 or less were considered significant. For animal studies, a sample size of five mice per group was found to be sufficient to have a high confidence of detecting a difference in means of at least 25% between groups (two-tailed test using *P* < 0.05) with a power of 90%, as determined by power analyses using the G*Power software (67). This number was consistent with numbers we have used in the past for similar studies and accounted for animal attrition due to mortality over the study period.

### Data availability

All data supporting the findings of this study are available within the manuscript, Supplementary Information, or the Source Data deposited in Hopkins Research Data Repository (Fluorescence imaging and Flow Cytometry metadata).

## Acknowledgments

The authors thank VEuPathDB for the readily accessible *T. gondii* genome assemblies and their functional annotations. We would also like to thank the Coppens lab for helpful suggestions regarding data analysis. We thank the individuals who provided plasmids, cell lines and parasite strains. We are also grateful to the EM Facility at the Johns Hopkins University (Michael Delannoy and Barbara Smith) for preparing EM samples. Light microscopy images were generated using instruments and support of the Light Microscopy Core of the Department of Molecular Microbiology and Immunology at the Johns Hopkins Bloomberg School of Public Health. This study was supported by the National Institute of Health grants R01AI166921 and R01AI138714 (IC). SR01AI138714 (IC), and [in part] by the Intramural Research Program of the National Institutes of Health (NIH). The contribution of the NIH author(s) are considered Works of the United States Government. The findings and conclusions presented in this paper are those of the authors(s) and do not necessarily reflect the views of the NIH or the U.S. Department of Health and Human Services. SMK recognizes pilot funding support from the Sherrilyn and Ken Fisher Center for Environmental Infectious Diseases at Johns Hopkins University School of Medicine, as well as the Johns Hopkins Malaria Research Institute. The funders had no role in study design, data collection and analysis, decision to publish, or preparation of the manuscript.

## Author contributions

SMK and IC conceived the ideas; IC designed the experiments in vitro; SMK designed virulence and immunization experiments; VP, KE and MEG designed DGAT1 constructs and generated ΔDGAT1; SMK, IC, YFG, JDR, KE, VP, KSM, YZ and DW performed the experiments; SMK, IC, YFG, FZ, JDR, JS and MA analyzed data; SMK wrote the initial draft; IC, MEG, JS, MA, JDR, YZ and FZ revised the manuscript and provided editorial comments. All authors reviewed and approved the manuscript for publication. Contributions of Mustafa Akkoyunlu and Jiro Sakai are informal communications and represent their own best judgement. These comments do not bind or obligate FDA.

**Supplementary Fig. 1.**
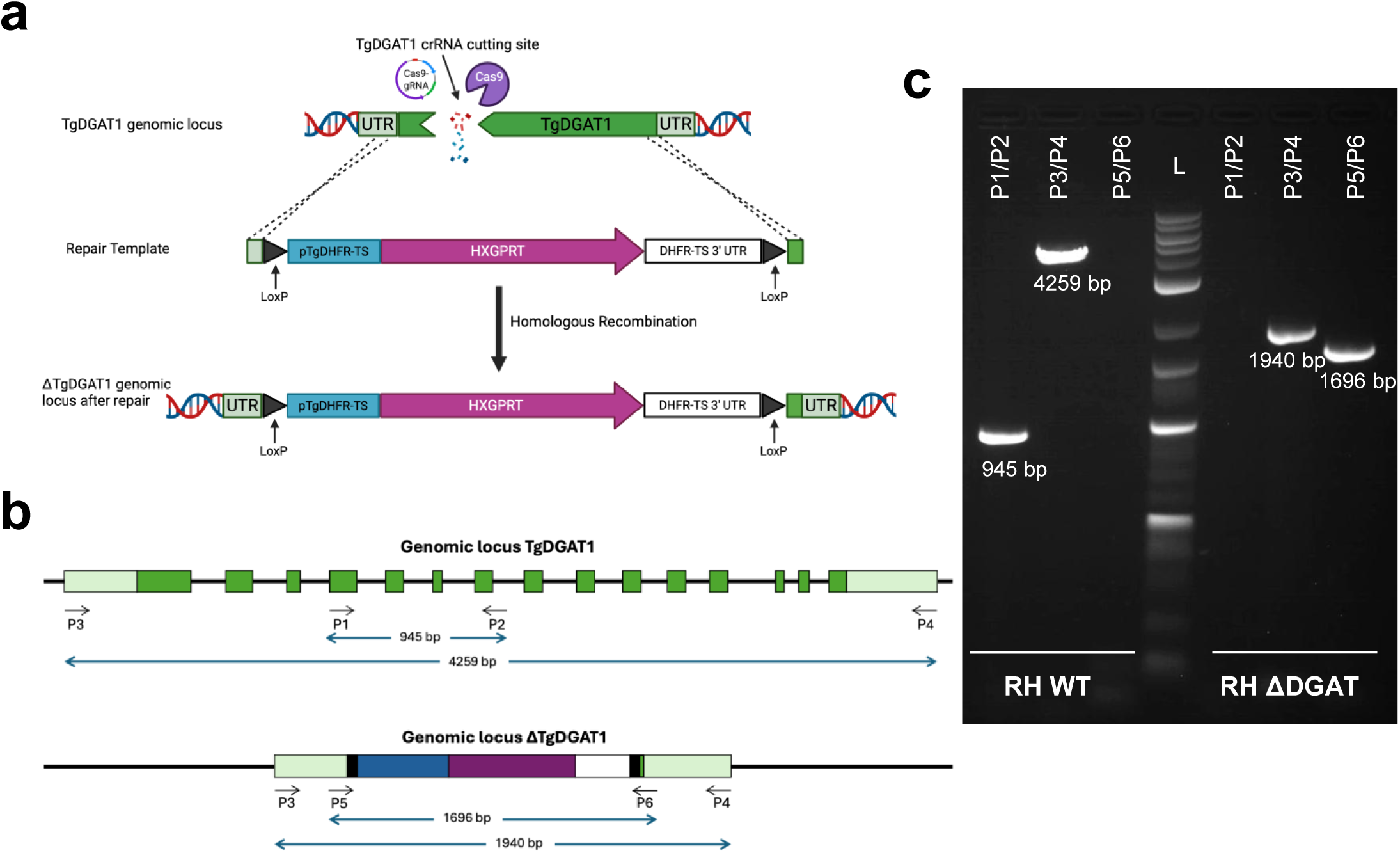
Creation of the *T*. *gondii* RH ΔDGAT1strain. (a) Diagram illustrating the strategy to disrupt TgDGAT1 (TGGT1_232730) in the type I *T*. *gondii* strain using CRISPR/Cas9 gene editing. The location of the guide RNA sequence is shown, indicating the predicted Cas9-induced DNA break in the first exon of TgDGAT1. The repair construct was designed to replace the TgDGAT1 gene with the HXGPRT selection cassette. (b) Diagram demonstrating the PCR screening method to identify *T*. *gondii* clones with the DGAT1 deletion. One primer pair (P1/P2) is located within the deletion, while another pair (P3/P4) is outside it. The third pair (P5/P6) detects correct integration of the repair template. (c) Representative PCR results from transfected *T*. *gondii* parasites confirming the deletion of the DGAT1 gene at the genomic level and replacement with the HXGPRT marker. DNA from both parental (WT) and transfected clones was used as a template. Unlike the WT parasites, the ΔDGAT1 clone lacks the P1/P2 band, shows a smaller P3/P4 band, and has a P5/P6 band, indicating successful deletion and correct marker integration. L indicates an 1-kb plus DNA ladder.

**Supplementary Fig. 2.**
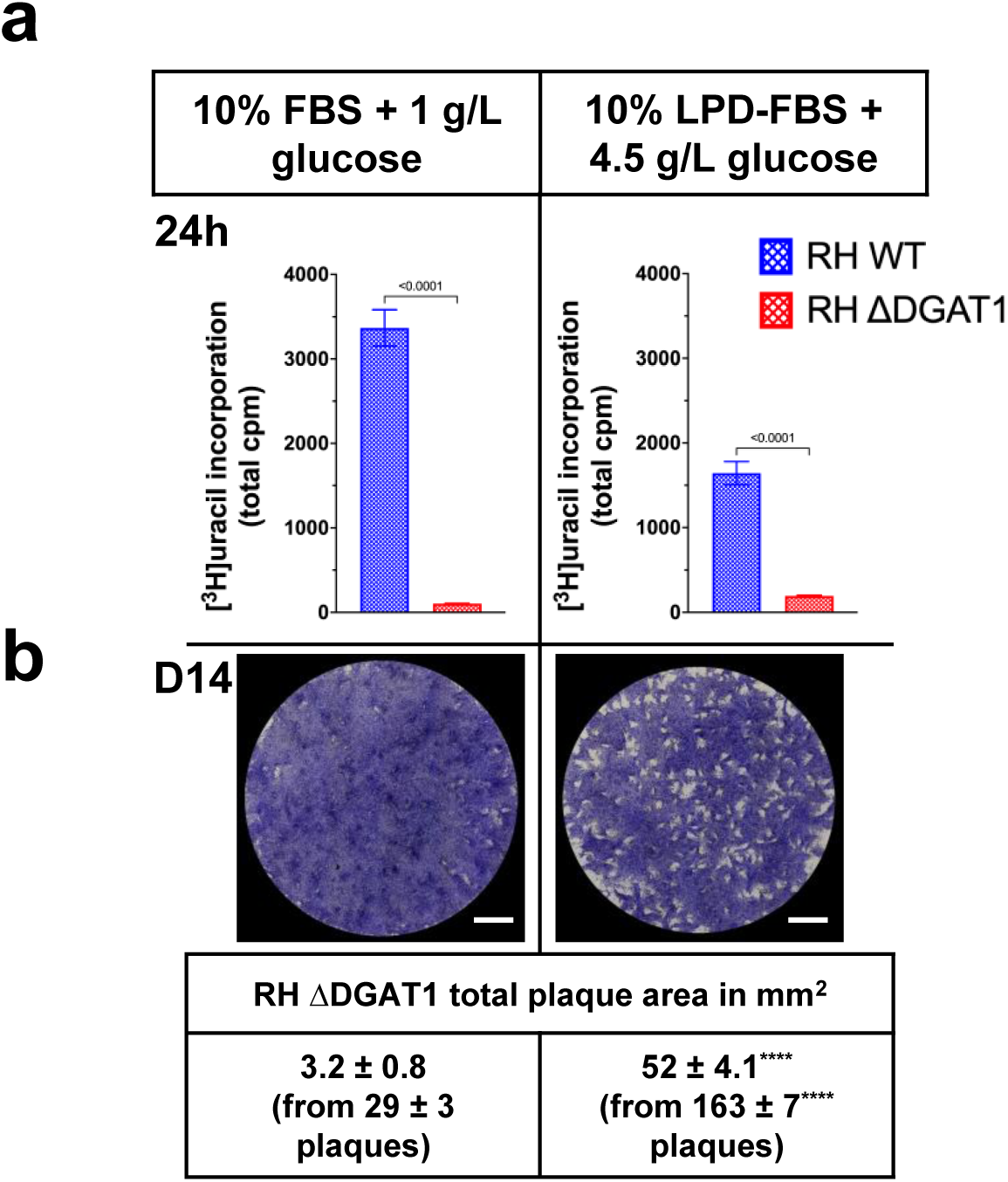
*Toxoplasma* RH ΔDGAT1 strain shows improved growth in low-lipid and high-glucose conditions. The RH WT or RH ΔDGAT1 *T*. *gondii* strain was cultured in either standard MEM with 10% FBS and 1 g/L glucose or in modified MEM with 10% LPD-FBS and 4.5 g/L glucose. (a) Parasite replication in human fibroblasts was quantified by [^3^H]-uracil incorporation at 24 hours post-infection, with comparisons between standard and modified media. Results show a slight recovery in replication defects in the absence of lipids and in the presence of added glucose. HFF-1 monolayers grown in both media were infected with either the WT or ΔDGAT1 strain for 24 hours, followed by incubation with tritiated uracil. Data expressed as mean ± SD from 2 independent assays were analyzed using an unpaired two-tailed t-test. (b) Analysis of parasite growth was conducted using plaque assays on fibroblast monolayers after 14 days of infection with 200 RH ΔDGAT1 parasites to compare growth in standard and modified MEM. Mutant parasites showed significantly better growth in the modified medium. Representative images and quantification of the total lysed area and number of plaques (means ± SD) from 3 independent experiments are shown; ****, *p* < 0.0001 (unpaired two-tailed t-test). All scale bars, 5.25 mm.

**Supplementary Fig. 3.**
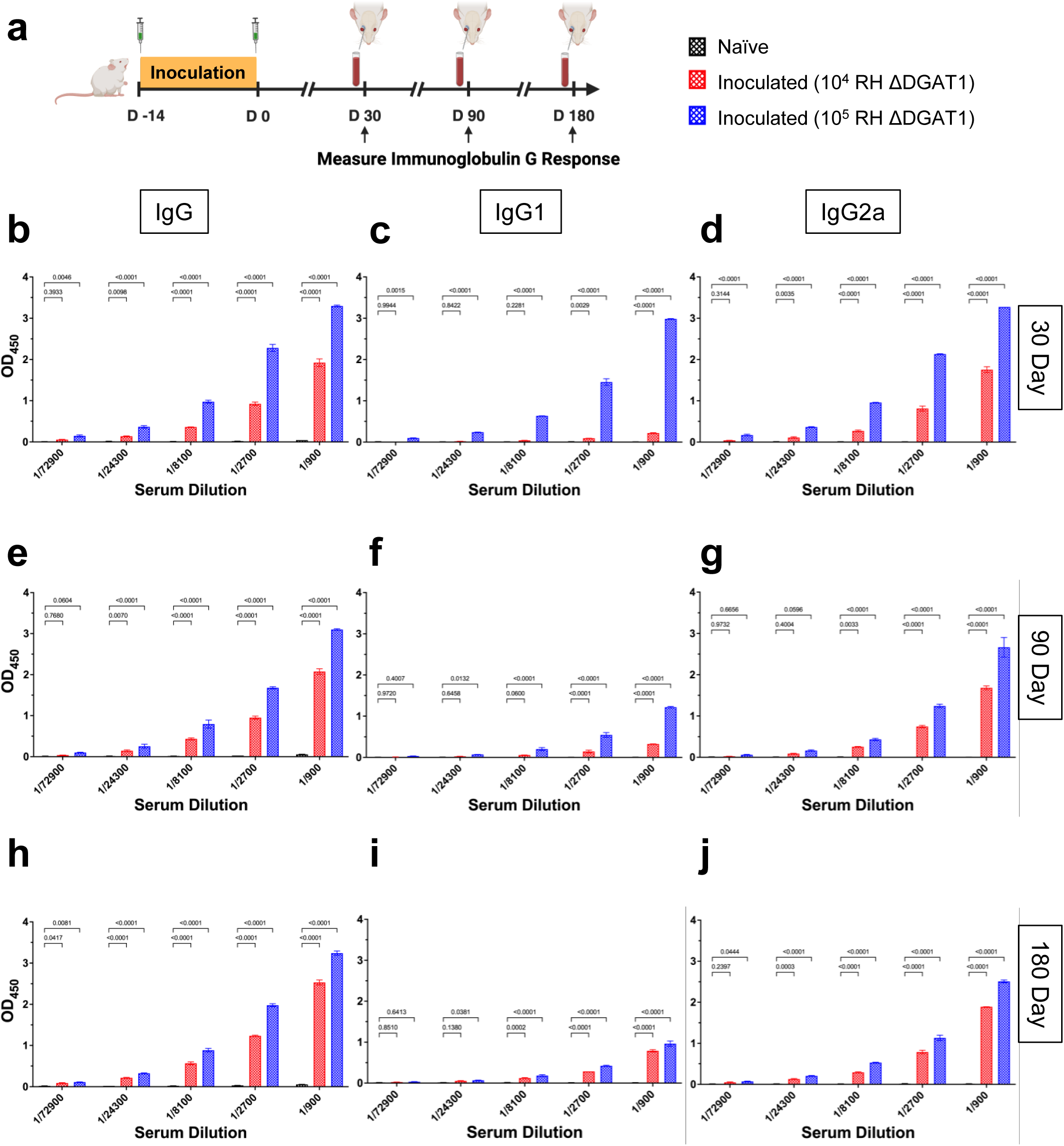
Inoculation of the ΔDGAT1 mutant induces strong antibody responses against *T*. *gondii* in mice. (a) Timeline for detection of *T. gondii*-specific IgG response in BALB/c mice by ELISA. Mice (n = 5 per group) were inoculated with two doses of 10^4^ or 10^5^ RH ΔDGAT1 parasites, and IgG, IgG1, and IgG2a levels were measured in the serum of naïve and vaccinated mice at 30, 90, and 180 days post-inoculation. Relative levels of *Toxoplasma*-specific IgG were determined by indirect ELISA using immuno-plates coated with 10 μg/mL of soluble *T. gondii* antigens. (b-j) Data for serum dilutions ranging from 1/900 to 1/72900 are shown. Mice immunized with RH ΔDGAT1 showed significantly higher levels of IgG (b, 30 days; e, 90 days; h, 180 days) and its subclasses IgG1 (c, 30 days; f, 90 days; i, 180 days) and IgG2a (d, 30 days; g, 90 days; j, 180 days) than the control group. Statistical analyses were performed using two-way ANOVA with Dunnett’s multiple-comparison test. Mean ± SD data with *p*-values are shown.

**Supplementary Fig. 4.**
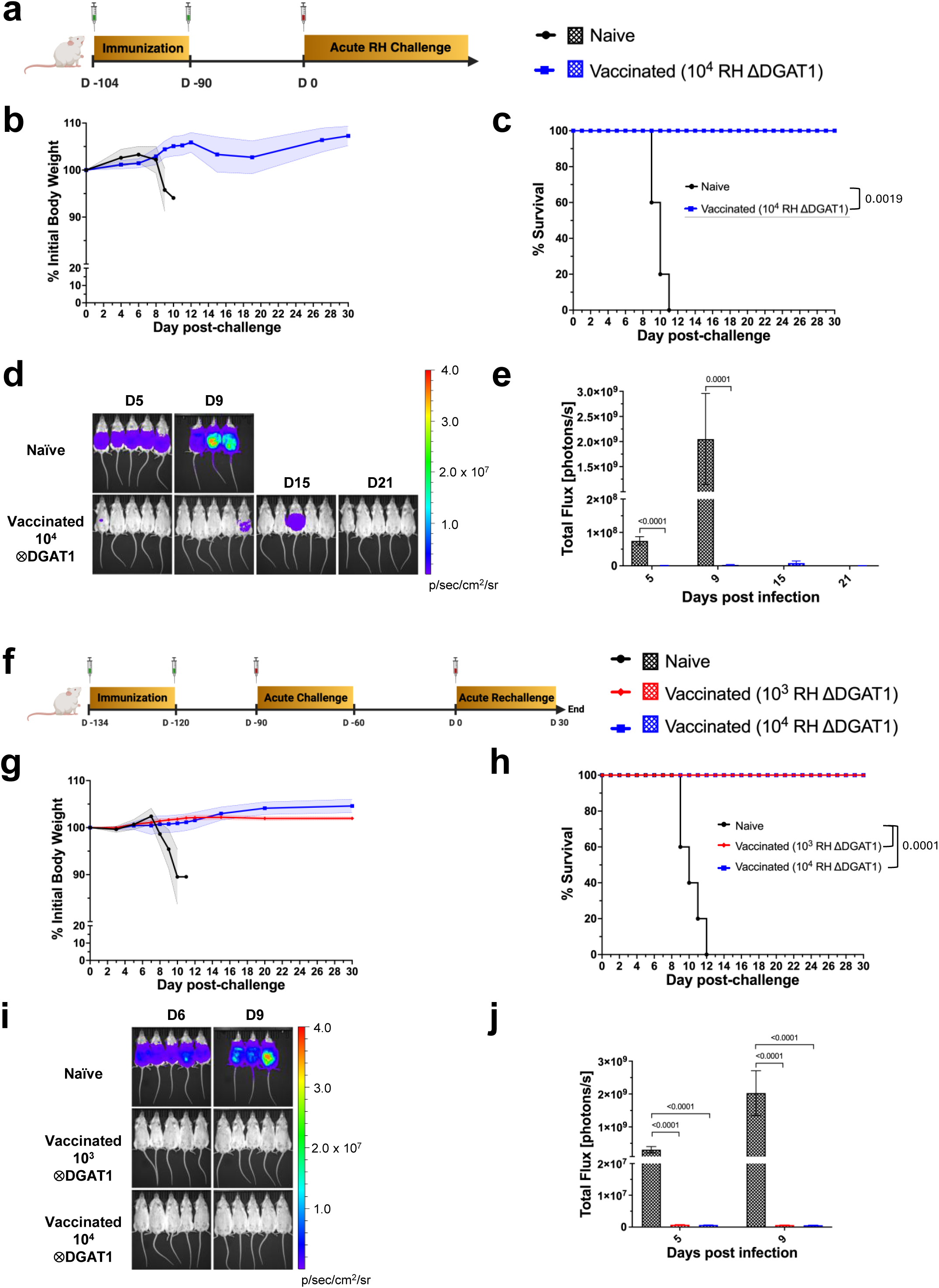
Vaccination with RH ΔDGAT1 parasites protects mice against acute type I *T. gondii* infection. (a) Female CD-1 mice (n = 5 per group) received two immunizations of 10^4^ RH ΔDGAT1 parasites, two weeks apart, followed by challenge with 500 tachyzoites of the RH luciferase *T*. *gondii* strain 90 days post-final immunization. (b) Normalized weights (means ± SEM) and (c) survival of naïve and immunized mice were plotted. Vaccinated mice showed no signs of infection or weight loss over 30 days, while naïve mice lost weight rapidly and died quickly by day 11. The log-rank (Mantel–Cox) test assessed survival differences. (d) Bioluminescence images of naïve and immunized mice at days 5, 9, 15, and 21 post-infection. Vaccinated mice exhibited no luminescence at day 21. (e) Whole-body luminescence quantification (mean ± SEM, photons/sec), with *p*-values from multiple unpaired t-tests with Holm-Sidak’s correction. (f) Female CD-1 mice (n = 5 per group) were immunized twice with either 10^3^ or 10^4^ RH ΔDGAT1 parasites. Thirty days after the last immunization, they were challenged with 500 tachyzoites of the type I RH luciferase-expressing *T*. *gondii* strain. All vaccinated mice survived the initial challenge and were later rechallenged 90 days afterward. (g) Normalized weights (means ± SEM) and (h) survival for naïve and immunized mice were plotted. Vaccinated mice displayed no infection symptoms or weight loss during 30 days, while naïve mice quickly lost weight and died within 12 days of infection. The log-rank (Mantel–Cox) test was used to assess the statistical significance of the difference in survival probability between naïve and vaccinated mice. (i) Bioluminescence images of naïve and immunized mice are shown at days 6 and 9 post-infection with luciferase-expressing RH parasites. Vaccinated mice exhibited no detectable parasite luminescence. (j) Quantification of the total body luminescence from luciferase-expressing parasites in CD-1 mice, shown in (i). Mean ± SEM data (total flux, photons/sec) were plotted, and *p*-values were determined using multiple unpaired t-tests with Holm-Sidak’s correction for multiple comparisons.

**Supplementary Fig. 5.**
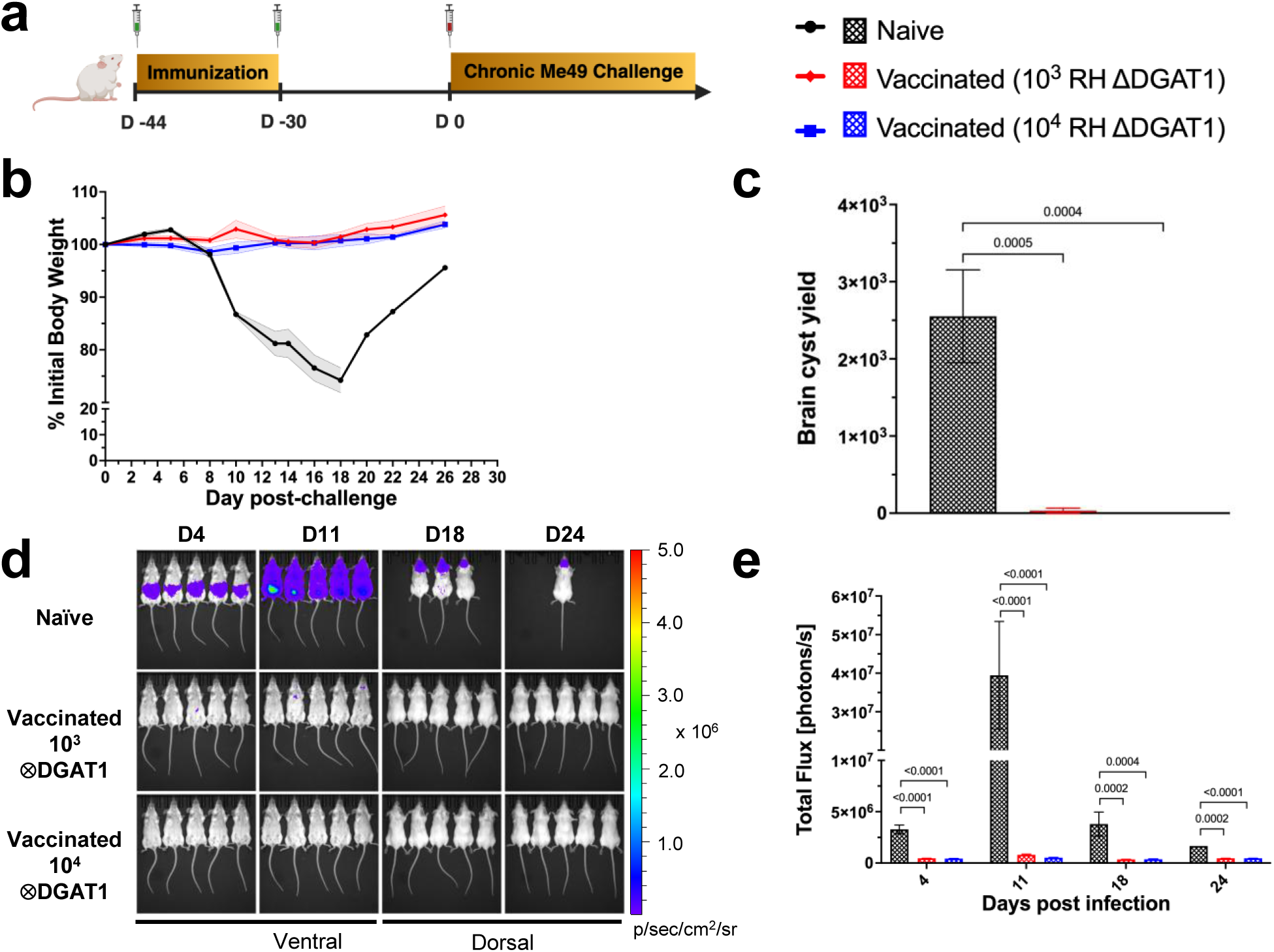
Immunization of mice with RH ΔDGAT1 parasites prevents cyst formation due to type II *T*. *gondii* infection. (a) Female BALB/c mice (n = 5 per group) were given two immunization doses of 10^3^ or 10^4^ RH ΔDGAT1 parasites, spaced two weeks apart, followed by challenge with 2000 tachyzoites of the type II Me49 luciferase-expressing *T. gondii* strain 30 days after the final immunization. (b) Normalized weights (means ± SEM) for naïve and immunized mice were plotted for all surviving mice at the specified time. (c) Brain cyst numbers of surviving or moribund mice, euthanized at 30 days or after 2 weeks of infection, respectively, were determined using DBA staining. Mean ± SEM data are shown (*p*-values calculated by one-way ANOVA with Dunnett’s multiple-comparison test). (d) Bioluminescence images of naïve and immunized mice are shown on days 4, 11, 18, and 24 after infection with luciferase-expressing Me49 parasites. To assess brain cyst load in chronic infection, mice were imaged dorsally on days 18 and 24. (e) Quantification of whole-body luminescence of Me49-Luc parasites in BALB/c mice, shown in (d). Mean ± SEM data (total flux, photon/sec) were plotted, and *p*-values were obtained using two-way ANOVA with Dunnett’s multiple-comparison test.

**Supplementary Fig. 6.**
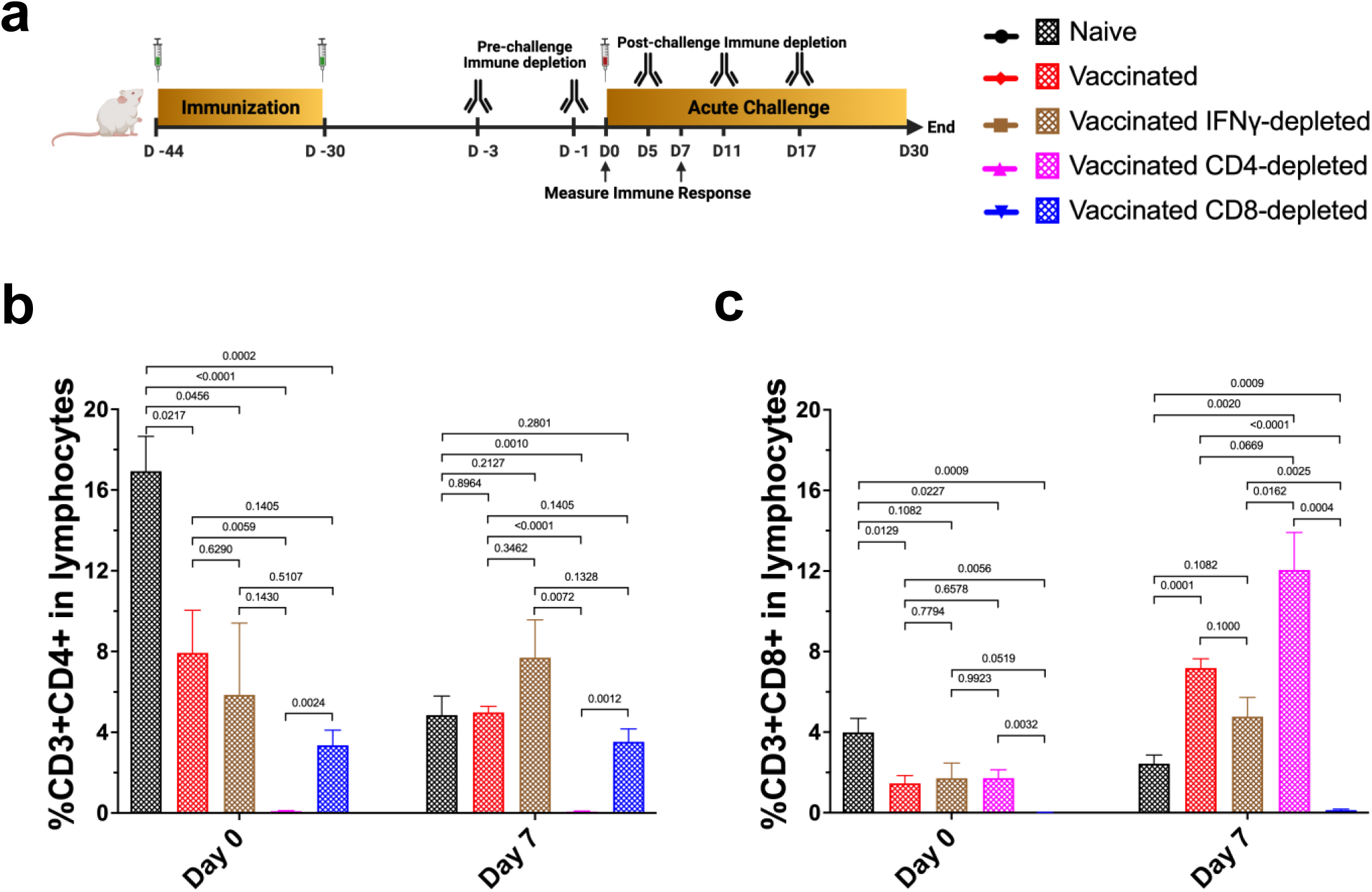
The immune response in mice following ΔDGAT1 mutant vaccination involves CD8^+^ T cells. (a) Female BALB/c mice received two doses of 10^4^ RH ΔDGAT1 parasites, two weeks apart. IFN-γ levels, CD4^+^ T cells, or CD8^+^ T cells were then depleted via intraperitoneal injections of 200 μg of anti-IFN-γ (Clone XMG1.2), anti-CD4 (Clone GK 1.5), and anti-CD8b (Clone H35-17.2) monoclonal antibodies, or mock isotype controls (Clones HRPN IgG1 and LTF2 IgG2b) on specified days, before and after challenge with a lethal dose of 500 luciferase-expressing type I RH tachyzoites. T-cell responses were measured in the serum of naïve and vaccinated mice on day 0 and day 7 of challenge. Percentages of (b) CD3^+^ CD4^+^ T cells and (c) CD3^+^ CD8^+^ T cells within peripheral lymphocytes of mice (n = 5 per group) were detected with flow cytometry. Results confirmed the successful depletion of CD4^+^ and CD8^+^ T cells in antibody-treated mice before and after infection. Data are presented as means ± SD. Statistical significance was determined using multiple unpaired t-tests with Holm-Sidak’s multiple-comparison test.

